# Inferring somatic mutation dynamics from genomic variation across branches within long-lived tropical trees

**DOI:** 10.64898/2026.04.02.716038

**Authors:** Sou Tomimoto, Akiko Satake

**Affiliations:** Department of Biology, Faculty of Science, Kyushu University, Fukuoka, Japan; Graduate School of Systems Life Sciences, Kyushu University, Fukuoka, Japan; Center for Frontier Research, National Institute of Genetics, Mishima, Japan

**Keywords:** Approximate Bayesian Computation (ABC), Dipterocarpaceae, Genetic mosaicism, Shoot apical meristem, Somatic genetic drift, Somatic mutation rate, Stem cell lineage, Subindividual variation

## Abstract

Trees accumulate somatic mutations throughout their long lifespan, resulting in genetic mosaicism among branches. While recent genomic studies quantified these mutations, they were largely limited to describing static patterns of variation. In this study, we developed a mathematical model to infer the dynamic processes of somatic mutation accumulation from snapshot genomic data obtained from four tropical trees (Dipterocarpaceae), which dominate tropical rain forests in Southeast Asia. Our model focus on genetic differences between shoot apical meristems (SAMs) at branch tips and explicitly incorporate stem cell dynamics within SAMs during shoot elongation and branching, enabling us to quantify somatic genetic drift arising from stem cell lineage replacement. By comparing model predictions with empirical data from Dipterocarpaceae trees, we estimated key parameters governing stem cell dynamics and somatic mutation rates. Our results indicate that both shoot elongation and branching involve replacement of stem cell lineages, leading to a moderate degree of somatic genetic drift. Accounting for stem cell dynamics resulted in slightly lower mutation rate estimates than previous approaches that ignored these processes. Using the estimated parameters, we further performed stochastic simulations to predict patterns of somatic mutations, including features not directly observed in the sampled trees, such as occasional deviations of somatic mutation phylogenies from physical architecture. Together, our modeling framework provides insights into how genetic mosaicism is shaped within tropical trees and reveals the stem cell dynamics underlying their long-term growth and accumulation of somatic mutations. (236 words)

**Highlights:**

- We built mathematical models to predict the genetic differences between branch tips by somatic mutations.
- The model considers the varying dynamics of stem cells in shoot meristem during shoot elongation and branching.
- We compared the model prediction with empirical data from tropical trees, Dipterocarpaceae, and estimated the dynamics of stem cells and mutation rate.
- Somatic mutation dynamics were shaped by somatic genetic drift arising from stem cell lineage replacement during shoot elongation and branching.
- Accounting for stem cell dynamics led to slightly smaller estimates of mutation rates compared with previous estimates that ignored the dynamics.
- Our models offer insights into how genetic variability is shaped in the tropical trees and the stem cell dynamics underlying their long-term growth.

## INTRODUCTION

Recent advances in sequencing technology have enabled the quantitative identification of somatic mutations accumulating in plants (Schoen & Schultz, 2019; Reusch *et al*., 2021). Accumulation of single nucleotide variants (SNVs) within individuals have been reported in many plant species, such as long-lived trees (Schmid-Siegert *et al*., 2017; Plomion *et al*., 2018; Wang *et al*., 2019; Hanlon *et al*., 2019; Orr *et al*., 2020; Hofmeister *et al*., 2020; Perez-Roman *et al*., 2022; Duan *et al*., 2022; Schmitt *et al*., 2024; Goel *et al*., 2024; Satake *et al*., 2024), cultivars (Wang *et al*., 2019; Adamek *et al*., 2022; Shirasawa *et al*., 2024), and clonal plants (Yu *et al*., 2020, 2024; Zheng *et al*., 2022). Somatic SNVs accumulating in pluripotent stem cells of the shoot apical meristem (SAM) are transmitted to newly formed shoots (Schmid-Siegert *et al*., 2017; Yu *et al*., 2020), shaping genetic structures that reflect an individual’s ontogeny (Schmid-Siegert *et al*., 2017; Perez-Roman *et al*., 2022; Goel *et al*., 2024; Satake *et al*., 2024). Somatic mutations appear largely neutral within trees (Orr *et al*., 2020; Perez-Roman *et al*., 2022; Duan *et al*., 2022; Satake *et al*., 2024) but undergo purifying selection in clonal plants (Yu *et al*., 2020; Zheng *et al*., 2022). These mutations can also be transmitted to offspring (Plomion *et al*., 2018; Wang *et al*., 2019; Goel *et al*., 2024).

Despite these advances, the empirical studies have been limited to capturing a snapshot of somatic mutations at specific stages of growth, leaving the dynamic processes by which these mutations accumulate and spread among branches during plant growth largely unknown. As somatic mutations accumulate in stem cells of SAMs, their dynamics, a process referred to as somatic genetic drift, affects the patterns of mutation accumulation (Yu *et al*., 2020; Reusch *et al*., 2021). For instance, topological congruence between physical architecture and phylogenetic tree derived from somatic mutations have yielded non-consistent results (Orr *et al*., 2020; Zahradníková *et al*., 2020; Schmitt *et al*., 2024; Goel *et al*., 2024; Satake *et al*., 2024). Although such inconsistencies may reflect differences in underlying stem cell dynamics (Zahradníková *et al*., 2020; Tomimoto & Satake, 2024), detailed dynamics cannot be inferred solely from snapshot genomic data.

Mathematical models have facilitated understanding of such dynamics of neutral somatic mutations. Early theoretical studies demonstrated that the fixation or loss of a mutated cell depends on the number of stem cells in a meristem and the rate of stem cell renewal through division (Klekowski & Kazarinova-Fukshansky, 1984a; Klekowski *et al*., 1989). A smaller number of stem cells and more frequent cell divisions lead to stronger somatic drift in the SAM during growth. With the growing availability of somatic mutation data from individuals of long-lived plants, methodological advances have started to bridge the gap between theoretical predictions and empirical observations. Tomimoto & Satake (2023) proposed a systematic modelling framework whose predictions can be directly compared with the observed patterns of somatic mutations within individuals, while accounting for diverse stem cell dynamics. They applied the model to the snapshot genomic data obtained from poplar trees (Hofmeister et al., 2020) and demonstrated that patterns of somatic mutation accumulation can differ significantly depending on the underlying stem cell dynamics, even under identical branching architectures and mutation rates. While this modelling approach successfully bridged snapshot data to stem cell dynamics, previous inference was hampered by a limited number of SNVs, necessitating strong assumptions about key parameters.

Here, we leverage a massive genomic dataset obtained from leaves of four tropical trees (Satake *et al*., 2024). These four individuals, representing two closely related Dipterocarpaceae species estimated to be 50–300 years old, provide independent biological replicates that enable us to infer the stochastic principles of mutation accumulation. Critically, the characteristics of this dataset require a modelling framework that can infer mutation dynamics from leaf-based genomic data without relying on stringent data requirements such as high-depth bud sequencing or histological observations. For example, Johannes (2025) inferred somatic dynamics in the L1 layer of apricot focusing on drift in branching (Goel *et al*., 2024), but this approach requires ultra-high-depth sequencing of bud samples, which is not available in the tropical trees. Similarly, Yu *et al*. (2024) calibrated a somatic genetic clock for clonal seagrass, but their approach relied on direct histological observations to constrain drift parameters, making it difficult to implement in the inaccessible canopies of long-lived tropical trees.

In this study, we extend the framework of Tomimoto & Satake (2023) for application to these tropical trees, incorporating somatic drift during both apical growth and branching into a unified Markov-chain framework. We estimated mutation rates, the number of stem cells, cell division rates, and stem cell dynamics governing somatic evolution. Based on these estimated parameters, we then conducted stochastic simulations to predict possible patterns of somatic mutations, including patterns not observed in the empirical data due to stochasticity. This modelling framework bridges snapshot patterns of intra-individual genomic variation and the stochastic dynamics of somatic mutation accumulation in long-lived tropical giants, which dominate tropical rain forests in Southeast Asia.

## MATERIAL AND METHODS

### Theoretical background

We first outline the model framework developed by Tomimoto & Satake (2023), which forms the basis for the extensions introduced in the present study.

Focusing on mutations accumulating in stem cells in a SAM, they modelled the plant growth as two processes: apical growth and branching (Fig. 1). Apical growth is the process of shoot elongation, during which stem cells repeatedly divide, resulting in the occurrence and accumulation of mutations in stem cell lineages (Fig. 1a). Branching is the process of new branch formation, in which the apical meristem of a main branch generates an axillary meristem that initiates a lateral branch (Fig. 1b). In branching, some stem cells in the apical meristem are stochastically sampled to form the axillary meristem, resulting in a bottleneck effect on somatic mutations. Thus, branch formation involves sampling a finite number of stem cell lineages, such that only mutations present in the sampled lineages are inherited by the new branch, giving rise to somatic drift through stochastic lineage dynamics. By iterating these processes, the model describes tree ontogeny and the accumulation of somatic mutations during growth (Fig. 1c).

**Figure 1.**
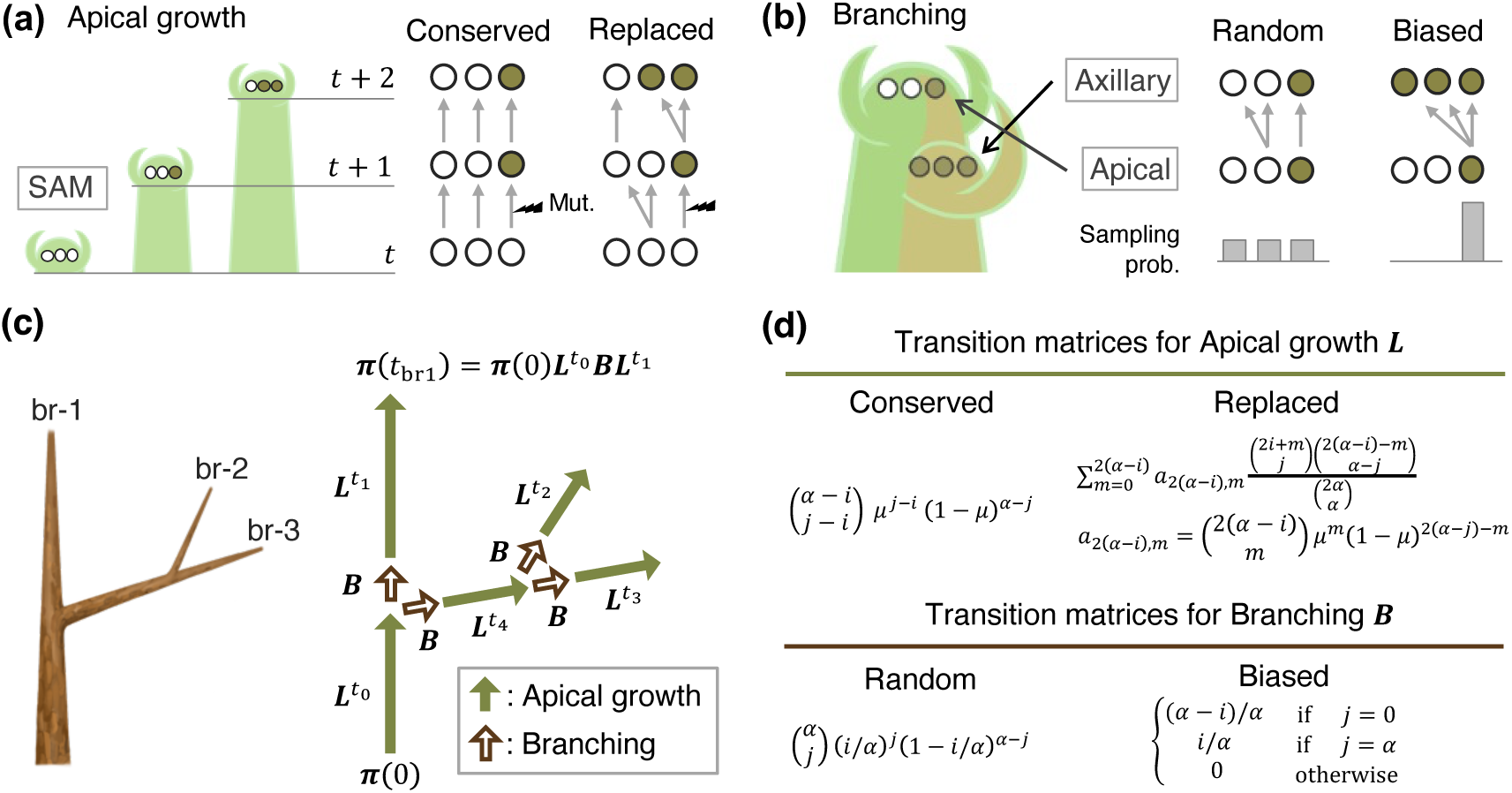
Model outline: somatic mutation dynamics in apical growth and branching. (a) Stem cell dynamics in apical growth (elongation). Stem cells (circles) undergo divisions and the shoot elongates, accumulating mutations. All lineages are retained in the shoot apical meristem in the conserved model, whereas stochastic lineage replacement occurs in the replaced model. (b) Stem cell dynamics in branching. Stem cells of an axillary meristem are sampled from the apical meristem. All stem cells are sampled with equal probability in the random model, whereas a single cell is sampled in the biased model. Green-filled circles indicate mutated cells, and open circles indicate non-mutated cells. (c) Translation of physical branching architecture (left) into a mathematical framework (right). The architecture is formed by repetitions of apical growth (green-filled arrow) and branching (brown-unfilled arrow). Mathematical symbols represent transition matrices for corresponding processes. For example, the mutational state of branch 1 (br-1) is given by 𝝅(0)𝑳t_0_ 𝑩𝑳t_1_, where 𝝅(0) indicates the initial state, 𝑡_0_and 𝑡_1_ are the number of cell divisions during apical growth for the corresponding branches. (d) Mathematical description of transition matrices: the 𝑖, 𝑗 elements for apical growth (𝑳) and branching (𝑩) are shown. Refer to SI Methods S1 for details.

To comprehensively describe varying degrees of somatic drift, Tomimoto & Satake (2023) considered contrasting stem cell dynamics in apical growth and branching. For apical growth, models without or with somatic drift were considered (Fig. 1a). In a conserved model (or structured model) without drift, asymmetric cell divisions occur successfully, and all stem cell lineages are conserved during apical growth. In a replaced model (or stochastic model) with drift, lineage replacement can occur by symmetric cell divisions. For branching, different strengths of the bottleneck effect were considered (Fig. 1b). In random branching (or unbiased branching), all stem cells in the apical meristem contribute to the formation of the axillary meristem with equal probability, allowing multiple cell lineages to be sampled. In biased branching, only a subset of stem cells contributes to the formation of the axillary meristem, resulting in a strong bottleneck effect (i.e., strong somatic drift). Their model accommodates diverse stem cell dynamics across plant species and can reproduce a wide range of somatic mutation accumulation patterns observed in the empirical studies (see SI Methods S1 for details).

However, their analysis relied primarily on computationally intensive stochastic simulations, which limited detailed fitting of the model to empirical data. We addressed this limitation by introducing a Markov chain–based mathematical model that extends their framework. This model mathematically drives the expected genetic differences between branch tips, substantially improving computational efficiency and enabling an exhaustive search of parameter space for model fitting. Using the estimated parameters, we then performed stochastic simulations to evaluate additional patterns, including somatic phylogeny, mutation distributions across branches, and mutation accumulation in SAMs.

### Outline of Markov chain model

The model describes the mutational state within a SAM, focusing on the number of mutated stem cells at a focal site (Klekowski *et al*., 1989; Tomimoto & Satake, 2023; SI Methods S1). For a SAM consisting of 𝛼 stem cells, the mutational state is represented by a probability vector 𝝅(𝑡) = (𝜋_0_(𝑡), 𝜋_1_(𝑡),…, 𝜋_a_(𝑡)), where 𝜋_i_(𝑡) denotes the probability that 𝑖 stem cells are mutated after 𝑡 growth events. At the beginning of plant development, no somatic mutations accumulate in a SAM with probability one, i.e., 𝝅(0) = (1, 0,…,0). From this initial state, somatic mutations accumulate in a SAM during growth. Changes in mutational state during apical growth and branching are described by transition matrices 𝑳 and 𝑩, respectively (Fig. 1c), assuming mutations occur independently across growth process. For instance, the state of a SAM that undergoes 𝑡_0_ apical growth events from the seedling stage is given by 𝝅(𝟎)𝑳to. If this SAM subsequently undergoes branching followed by an additional 𝑡_1_ apical growth, the state becomes 𝝅(𝟎)𝑳to 𝑩𝑳t1, which corresponds to the mutational state of branch br-1 in Fig. 1c.

This study newly introduces transition matrices for branching with random and biased sampling (Fig, 1c). By combining these with matrices for the conserved and replaced apical growth models (Klekowski *et al*., 1989; Tomimoto & Satake, 2023), we mathematically calculate of somatic mutation accumulation across tree ontogeny. The explicit forms of the transition matrices are given in Eqs. (S2–S5) in SI Methods S1.

### A measure for genetic differences between branch tip**s**

To compare model predictions with observations, we formulates the genetic difference between branch tips, or inter-branch SNVs, particularly considering the characteristics of leaf samples, as used in Satake *et al* (2024). A leaf originates from a subset of stem cells in a SAM (Poethig, 1989; Irish & Sussex, 1992), and mutations accumulating only in these stem cells are detected. A mutation shared by a larger proportion of stem cells is more likely to spread to leaves, as sectorial chimeras suggest (Stewart & Dermen, 1970; Ruth *et al*., 1985), and is more likely to be sampled and detected by sequencing. Thus, we assume that the probability of detecting a mutation is proportional to the frequency of the mutated stem cells within a SAM.

To reflect this frequency dependence, the measurement is defined by the average heterozygosity between cells randomly sampled from the SAMs of two different branches (Iwasa *et al*., 2024; Tomimoto *et al*., 2025). Let 𝜋*_i_^n^* be the probability of a branch 𝑛 having 𝑖 mutated stem cells in the SAM. The probability of sampling a mutated cell from the SAM is given by ∑_i_ 𝜋^n^𝑖/𝛼, where 𝑖/𝛼 is the probability of sampling one of 𝑖 mutated cells from 𝛼 stem cells. Then, the expectation of sampling a mutated stem cell from only one of the two branches 𝑛 and 𝑚 is given by: 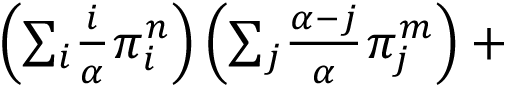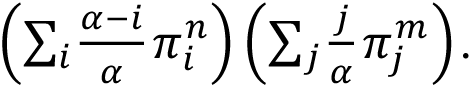 Here, the first term represents the probability of sampling a mutated cell from branch 𝑛 and a non-mutated cell from branch 𝑚. The second term represents the opposite case. Based on this definition, we derived the expectation of the inter-branch SNVs per nucleotide (refer to SI Methods S2 for full mathematical derivation). This measurement provides informative summary statistics for the analysis of leaf-sample genomic data, particularly when high-resolution SAM sequencing data is unavailable.

### Data of somatic mutations and branching architecture from tropical trees

We used published data on somatic single nucleotide variants (SNVs) detected from two tropical Dipterocarpaceae species, *Shorea laevis* and *Rubroshorea leprosula* (Satake *et al*., 2024). The dataset includes a substantial number of somatic SNVs identified from leaf samples collected at the tips of seven branches from four trees: 728 and 234 in two slow-growing *S. laevis* individuals (S1 and S2) and 106 and 68 in two fast-growing *R. leprosula* individuals (F1 and F2), respectively (Satake *et al*., 2024). Additionally, the branching architectures were recorded, allowing the model to simulate the growth history of trees.

Branching architectures, listed in Table S1 and Fig. S1, represent the growth history of apical growth and branching processes in the Dipterocarpaceae trees. We assume that apical growth occurs along a branch between bifurcation points, while branching occurs at bifurcation points. Apical growth events (stem cell renewal by cell division) are assumed to occur in proportion to branch length at a rate 𝑟, such that 𝑡 = 𝑟𝑑 apical growth events occur during a growth of length 𝑑 (Fig. 2a). This assumption reflects the fact that, in tropical trees lacking annual rings, branch length provides the most informative proxy for growth history. Branching events are assumed to occur symmetrically at bifurcation points for both resulting branches. This differs from Tomimoto & Satake (2023), who distinguished main and lateral branches, assuming that only lateral branches undergo the sampling process in branching. We adopt the present assumption because main and lateral branches are often morphologically similar and difficult to distinguish in mature tropical trees, e.g., supernumerary branch (Barthélémy & Caraglio, 2007).

**Figure 2.**
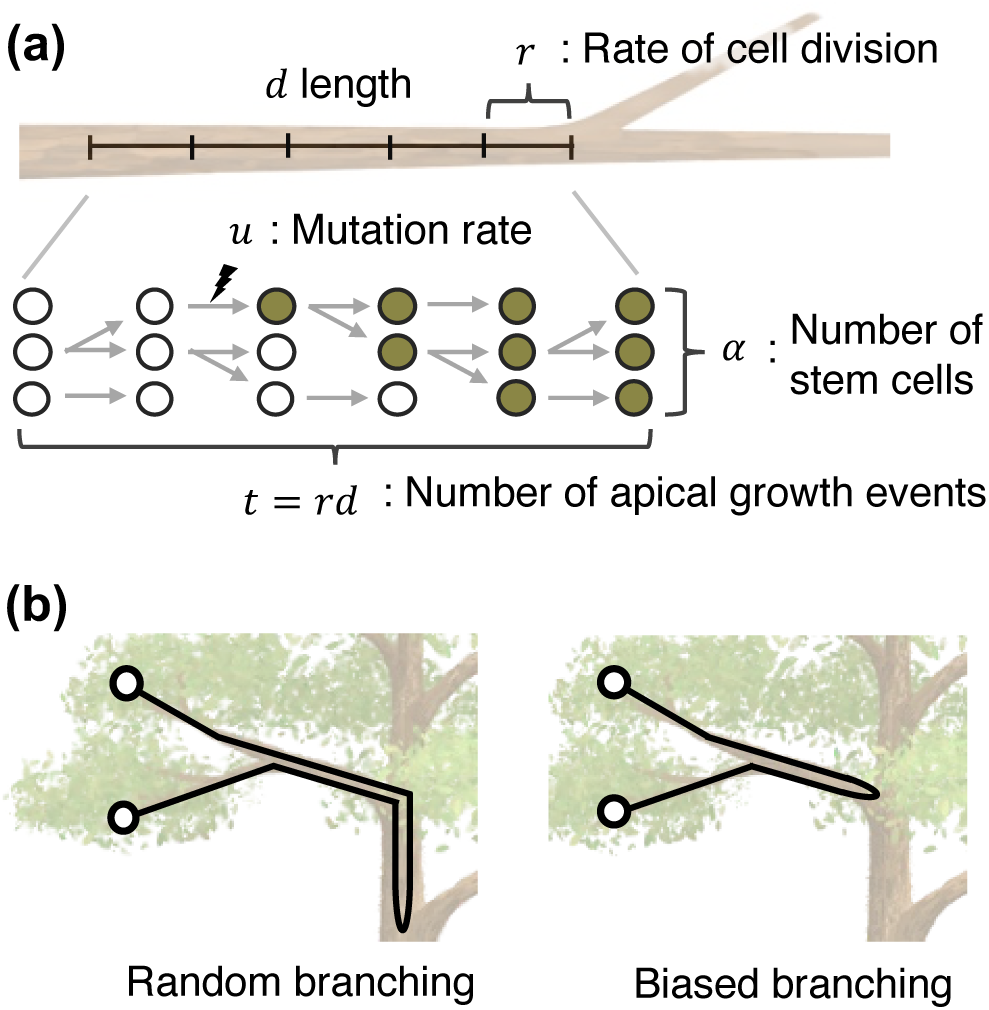
Factors affecting somatic genetic drift. (a) Model parameters. Somatic drift during apical growth is governed by the number of stem cells in a shoot apical meristem, 𝛼, and the rate of cell division, 𝑟. Each stem cell undergoes 𝑡 = 𝑟𝑑 divisions during 𝑑 length apical growth. Mutations occur at a rate 𝑢 per site per length. (b) Effects of sampling in branching. In contrast with random branching, biased branching leads to quick coalescence of stem cell lineages. Black lines represent lineage history of stem cells, and unfilled circles represent the focal stem cells at branch tips.

### Model fitting and parameter inference

To infer a model and parameters that fit the data, we applied approximate Bayesian computation with sequential Monte Carlo sampling (ABC-SMC) implemented in *pyABC* (Klinger *et al*., 2017; Schälte *et al*., 2022). ABC-SMC builds upon the standard ABC rejection algorithm. In this approach, parameter sets are sampled from prior distributions, and the resulting model predictions are compared with the observed data through a distance function. If the distance between prediction and observation is smaller than a threshold, the parameter set is accepted; otherwise, it is rejected. This process continues until the number of accepted parameter sets reaches a population size, forming one generation. The procedure proceeds over multiple generations with progressively tighter acceptance thresholds. Through this iterative refinement, the algorithm gradually focuses on regions of the parameter space that yield predictions closely matching the observations, thereby improving the posterior of parameters. In our analysis, we performed 25 generations with a population size of 300. The prior distributions of parameters are listed in Table 1.

**Table 1.** MAP estimates of parameters for each model.

| Symbol | Definition | Prior distribution | MAP for model I | MAP for model II | MAP for model III | MAP for model IV |
| --- | --- | --- | --- | --- | --- | --- |
| $\alpha$ | Number of stem cells | *Uniform* (1, 50) | 2.3804e+00 | 3.4186e+00 | 4.2525e+01 | 4.4291e+01 |
| $r$ | Number of cell divisions per meter apical growth | *Uniform* (1,33*) | 1.5534e+01 | 1.3628e+01 | 1.3429e+01 | 2.3872e+01 |
| $u_{S1}$ | Mutation rate per site per unit growth for S1 | *Uniform* (0,10<sup>-8</sup>) | 4.9444e-09 | 4.8152e-09 | 6.2577e-09 | 6.8651e-09 |
| $u_{S2}$ | Mutation rate per site per unit growth for S2 | *Uniform* (0,10<sup>-8</sup>) | 2.0834e-09 | 1.9326e-09 | 3.0020e-09 | 3.4863e-09 |
| $u_{F1}$ | Mutation rate per site per unit growth F1 | *Uniform* (0,10<sup>-8</sup>) | 6.9178e-10 | 8.5487e-10 | 1.1921e-09 | 1.3752e-09 |
| $u_{F2}$ | Mutation rate per site per unit growth F2 | *Uniform* (0,10<sup>-8</sup>) | 1.0088e-09 | 6.9577e-10 | 1.1110e-09 | 1.3273e-09 |

\* The maximum number of cell divisions per meter, 33, comes from the estimated value in *Arabidopsis thaliana* (Watson *et al.*, 2016).

As a distance function, we adopted the mean squared error (*MSE*) as follows: 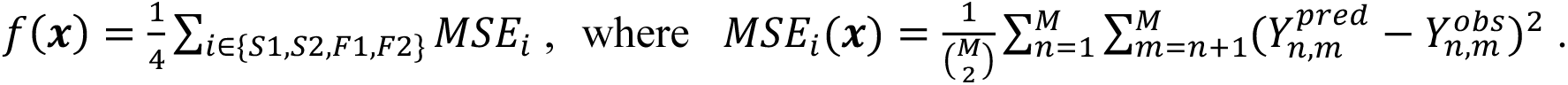 Here, 𝑖 ∈ {𝑆1, 𝑆2, 𝐹1, 𝐹2} denotes tree individuals. 𝑌*_n,m_^pred^* and 𝑌*_n,m_^obs^*: are the expected genetic difference between branch 𝑛 and 𝑚 from model prediction and sequence data, respectively. The denominator is the number of branch pairs (the number of branches in a tree is seven in our data, 𝑀 = 7).

For each model *j* ∈ {Conserved-random, Conserved-biased, Replaced-random, Replaced-biased}, we inferred the following parameter set: 𝒙_j_ = {𝛼, 𝑟, 𝑢_s1_, 𝑢_s2_, 𝑢_F1_, 𝑢_F2_}. Here, 𝛼 denotes the number of stem cells in a SAM, and 𝑟 is the rate of stem cell renewal per unit length growth during apical growth (i.e., cell division rate). The parameter 𝑢_i_ represents the mutation rate of an individual 𝑖 ∈ {𝑆1, 𝑆2, 𝐹1, 𝐹2}, which is scaled per unit length growth to allow direct compare with estimations from Satake *et al*. (2024). In the Markov chain model, the mutation rates were thus converted to per-division rates as 𝜇_i_ ≡ 𝑢_i_/𝑟. We assume that the model of stem cell dynamics, 𝛼, and 𝑟 are shared by *S. laevis* and *R. leprosula*, because their close phylogenetic relationship implies similar developmental processes (Ghazoul, 2016). In contrast, mutation rates are allowed to vary among individuals, reflecting the empirical observation that mutation rates also depend on individual age (Satake *et al*., 2024).

### Simulating the various patterns of somatic mutations

Using the MAP estimate of the parameters for each model of stem cell dynamics, we conducted stochastic simulations based on the model from Tomimoto & Satake (2023) and predicted various patterns of somatic SNVs within an individual. These patterns allow us to qualitatively evaluate the four models of stem cell dynamics from multiple perspective. Moreover, repeated stochastic simulations allow us to characterize the full spectrum of patterns that are probabilistically expected to arise, including those not observed in the empirical samples.

We evaluated the model predictions based on four somatic SNV patterns. First, we calculated the inter-branch SNVs and the probability distribution of *MSE* from them to check the estimation of model fitting. Second, we calculated phylogenetic trees from the simulated somatic SNVs by UPGMA and compared their topologies with the physical branching architectures by unweighted Robinson-Foulds distance (Robinson & Foulds, 1981). Third, we calculated the distribution patterns of somatic SNVs across branches, which is known to vary with the underlying stem cell dynamics (Tomimoto & Satake, 2023). Fourth, we calculated the number of accumulated SNVs in each SAM and their frequencies within the stem cell population.

We conducted 250 runs of simulation for each individual and stem cell dynamics (i.e., 250×4×4 simulations in total). Simulations used MAP estimates of parameters (Table 1). Other parameters and the branching architectures are listed in Table S2 and S1, respectively. For the details of simulation model, refer to Tomimoto & Satake (2023). Python codes for the simulations are available on https:#####.

## RESULTS

To provide a basis for interpreting observed empirical data, we first characterized how the models of stem cell dynamics and their associated parameters (𝛼, 𝑟, and 𝑢) shape the patterns of inter-branch SNVs (Fig. 3a,b). Inter-branch SNVs are primarily determined by the strength of somatic genetic drift during apical growth. Under strong somatic drift characterized by replaced apical growth, with a small number of stem cells 𝛼 and a high rate of cell divisions 𝑟 (Fig. 2a), inter-branch SNVs increase linearly from the origin, with steeper slopes at higher mutation rates 𝑢 (Fig. 3b). As somatic drift weakens, genetic heterogeneity among stem cells is preserved within the SAM. This heterogeneity maintained at the forking point leads to a positive intercept in the relationship between inter-branch SNVs and physical distance, i.e., mutations accumulated before forking contribute to genetic differences in addition to those arising after forking (Fig. 3b; Tomimoto *et al*., 2025). The magnitude of the intercept increases with weaker drift, i.e., a larger number of stem cells and a lower division rate, serving as an indicator of the strength of somatic drift (Figs. 2a, 3a).

**Figure 3.**
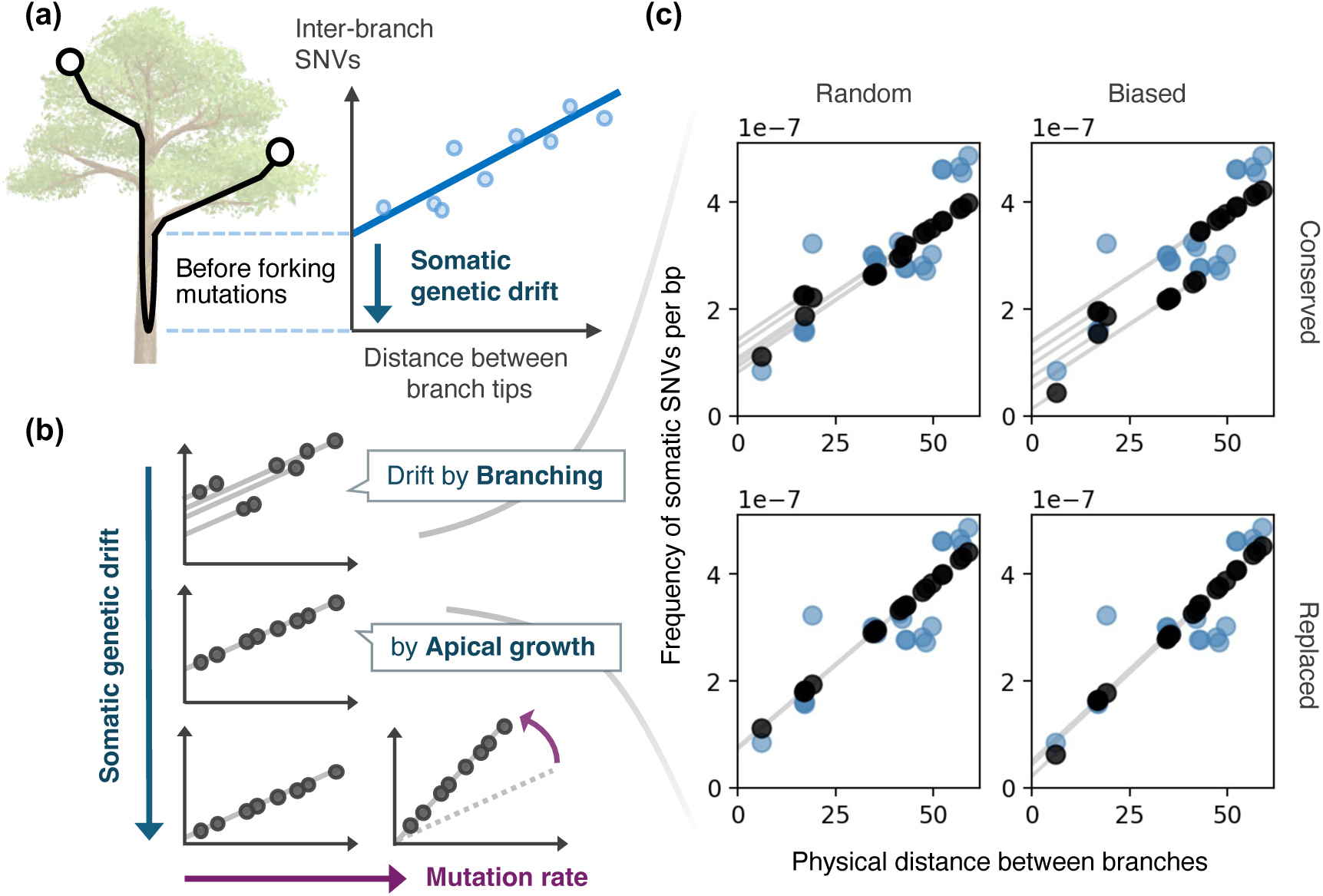
Inter-branch SNVs: conceptual framework and inferred patterns. (a) Genetic differences against the physical distance between branch tips. The intercept of this relationship quantifies the degree of somatic genetic drift. Weak somatic drift maintains genetic heterogeneity within the shoot apical meristem (SAM), which allows mutations accumulated before forking to contribute to inter-branch genetic differences, leading to a positive intercept. (b) Conversely, strong drift leads to rapid mutation fixation, causing the intercept approaching zero. The slope of this relationship is determined by mutation rate per unit length. (c) Predicted inter-branch SNVs by inferred models. Predictions for S1 individual from all models are plotted, consistently suggesting the positive intercepts. Black circles indicate model predictions, while blue circles represent observations from S1 individual. The parameters used for predictions come from MAP estimates for each model, which are listed in Table 1.

When somatic drift during apical growth is weak, branching becomes an important determinant of the pattern (Fig. 3b). Somatic drift by branching can lead to a smaller intercept than expected from apical growth alone. Strong drift by biased branching reduces genetic heterogeneity among stem cells and produces smaller intercepts (Fig. 2b), resulting in characteristic sawtooth patterns (Figs. 3b, S3). This effect is attenuated by random branching with weaker drift, as well as by a larger number of stem cells. Together, the slope, intercept, and fine-scale deviations in inter-branch SNV patterns emerge from the interplay between mutation rate and somatic drift (apical growth and branching dynamics, the number of stem cells, and cell division rate), as schematically summarized in Fig. 3, and provide the basis for model fitting.

### Inference of moderate somatic genetic drift in Dipterocarpaceae trees

By applying our models to the observed data from Dipterocarpaceae trees, we found that despite differing assumptions regarding stem cell dynamics, all models consistently suggest a moderate degree of somatic genetic drift. The optimal models, based on MAP estimates of parameters (Table 1; Fig. 4), successfully predicted the characteristic small intercepts observed in the empirical data, which signify moderate drift (Fig. 3c). This convergence on a consistent drift level arises through model-specific parameter combinations that compensate for their respective assumptions. For instance, the conserved models inferred a small stem cell number (𝛼 ≈ 2–3; Fig. 4b), attributing the necessary drift primarily to the branching process. In these models, a small 𝛼 is essential to generate the observed drift, as they assume no lineage turnover during apical growth (Figs. 4, S2–3A). Conversely, the replaced models inferred a substantially larger number of stem cells (𝛼 ≈ 40; Fig. 4b). Under the continuous lineage replacement during apical growth, the large size of stem cell population is required to maintain genetic heterogeneity within a SAM to provide the best fit to the moderate intercepts observed in Dipterocarpaceae trees (Figs. 3c, S4–5A). This robust identification of somatic drift, common to all models, ensures our estimation of how mutations accumulate over time.

**Figure 4.**
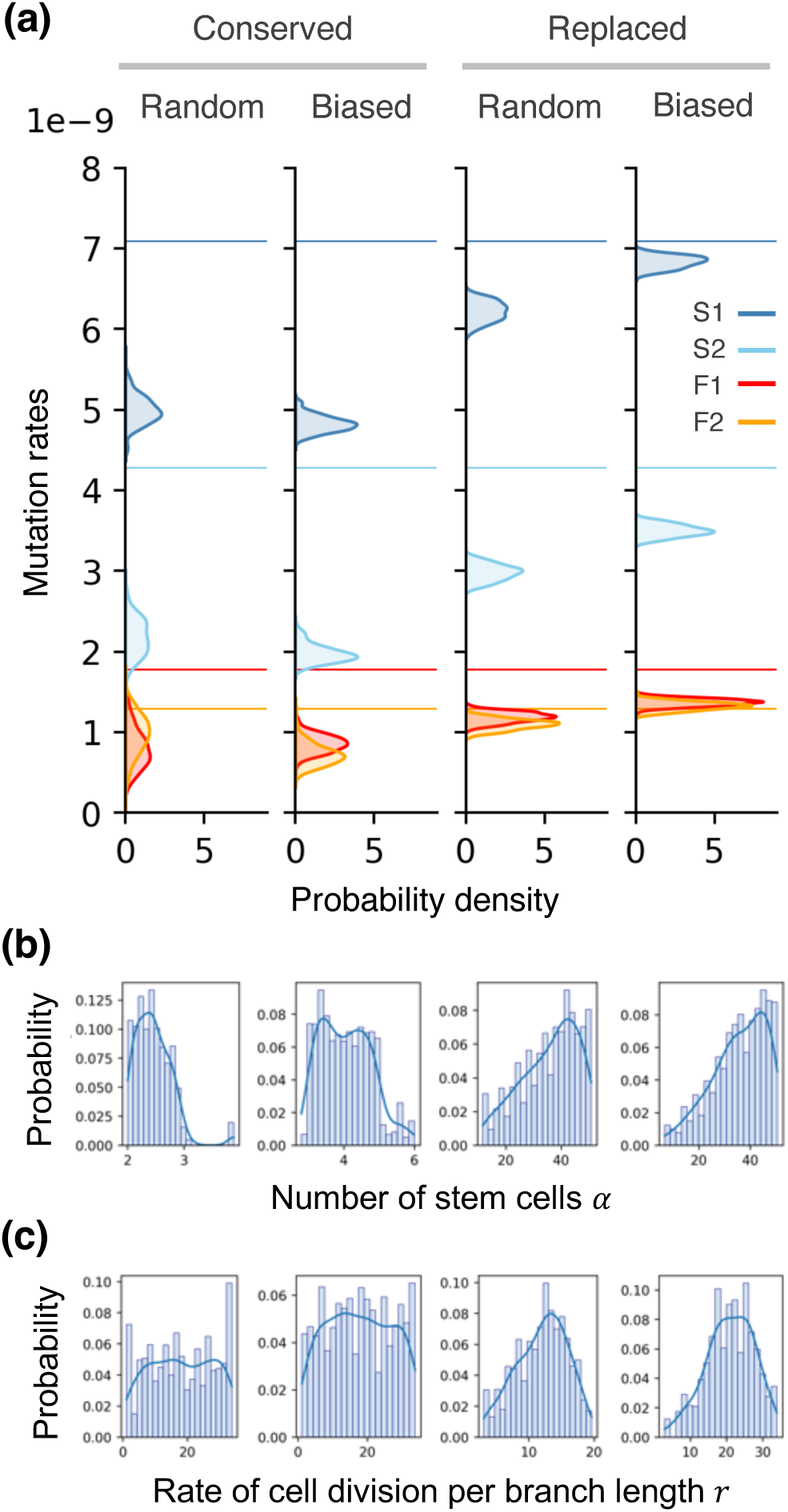
Posterior distributions of parameters estimated by model fitting. (a) Posterior distributions of mutation rates for each model. From left to right: conserved random, conserved biased, replaced random, replaced biased. Distributions are shown as kernel density estimations (KDEs). Horizontal lines indicate the rates estimated by Satake *et al*. (2024). Colors represent individuals (blue: S1, light blue: S2, red: F1, orange: F2). (b) Posterior distribution of the number of stem cells. (c) Posterior distributions of the rate of cell division per unit branch length. The order of models is the same as in panel A. Distribution curves are shown as KDEs.

Crucially, accounting for these stem cell dynamics led to a downward revision of estimated somatic mutation rate (Fig. 4a). Regardless of stem cell dynamics, all models consistently inferred mutation rates lower than those previously reported. This discrepancy arises because previous estimates, derived from linear regressions forced through the origin, did not account for the genetic heterogeneity accumulated prior to forking (i.e., the intercept). By ignoring this pre-existing variation, previous approaches overestimated the mutation rate. Although all models yielded clear posterior peaks for the mutation rate (Fig. 4a), we observed that the conserved models inferred lower rates than the replaced models. This difference reflects how each model partitions the observed inter-branch SNVs. The conserved models, characterized by weaker drift, attributed a larger proportion of genetic differences to mutations accumulated before forking, resulting in higher intercepts. In contrast, the replaced models placed greater weight on mutations arising after forking, resulting in a steeper slope by higher mutation rates (Fig. 4c). This raises a question: which of these contrasting models more accurately reflects the actual dynamics of somatic mutation accumulation in the tropical trees?

Although all models provided a reasonably good match to the observed data (Figs. 3c, S10, S11), the replaced models yielded superior predictions, suggesting that decades of apical growth in Dipterocarpaceae trees involves the continuous replacement of stem cell lineages. The minimum distances between predictions and observations, quantified by MSEs, were consistently smaller in the replaced models (Figs. S6–9A). This was further supported by stochastic simulations based on MAP estimates of parameters, where MSE distributions were consistently lower for the replaced models in all individuals (Fig. 5a). Beyond simple error metrics, the replaced models also better captured other patterns of mutation accumulation, such as topological congruence of the somatic phylogeny and the distribution of somatic SNVs. These findings provide compelling evidence that dynamic lineage turnover during apical growth, rather than strict conservation, governs the somatic SNV accumulation within Dipterocarpaceae trees, suggesting the slightly lower mutation rates than previous estimates.

**Figure 5.**
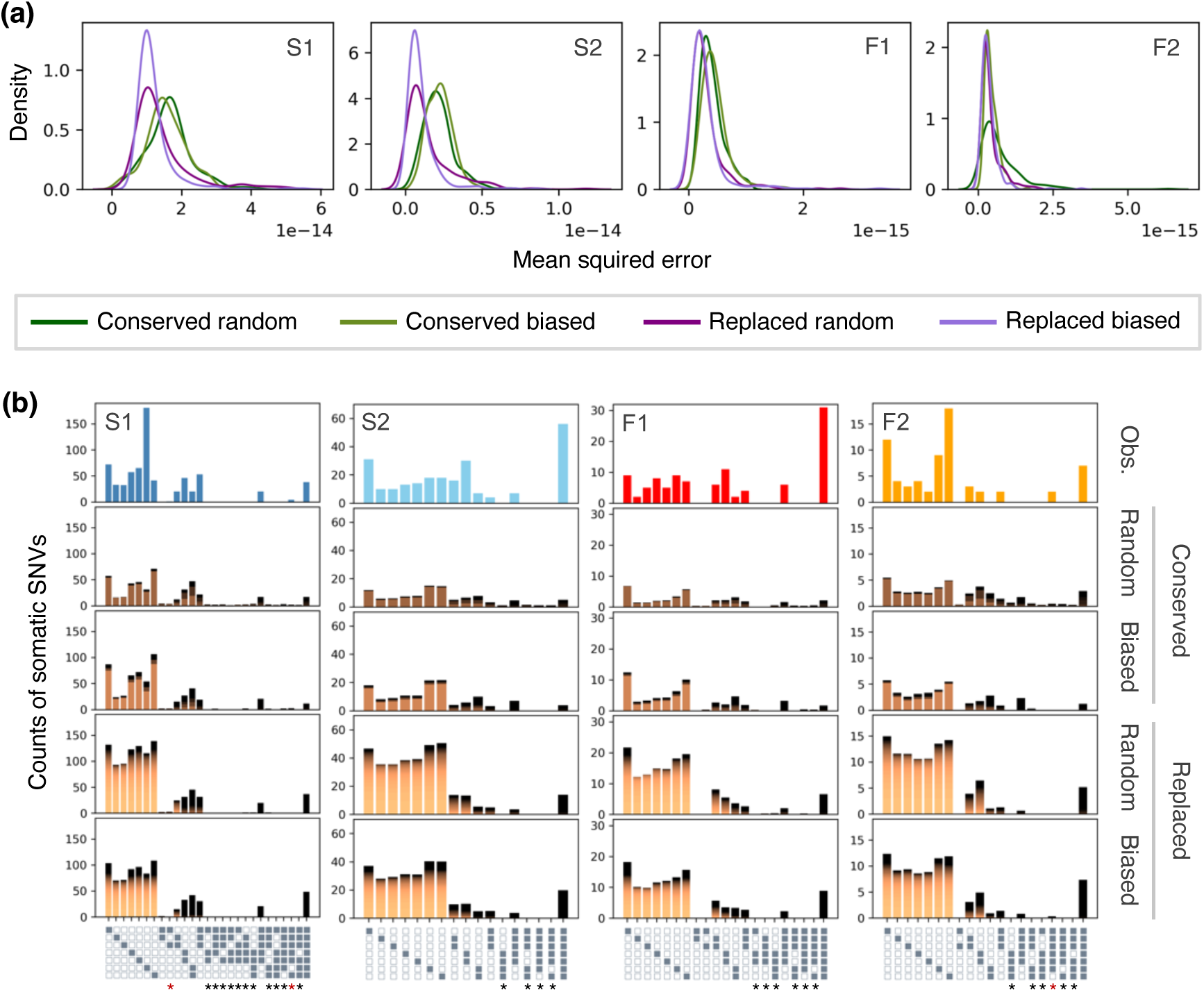
Somatic SNVs patterns predicted by simulations based on MAP estimates of parameters. (a) Probability distribution of mean squared errors between observation and prediction of inter-branch SNVs. Predictions come from 250 runs of stochastic simulations. The probability distributions were estimated using kernel density estimation. (b) Spatial distribution patterns of somatic SNVs across branches from the observations (top) and predictions (lower rows). Gray and white panels on the x-axis indicate the presence and absence patterns of a somatic SNV in each of the seven branch tips (from the top, corresponding branch IDs are 40, 30, 31, 20, 21, 10, 11). Asterisks indicate non-nested, patchy patterns; observed patterns are colored in red. Predictions are the average of 250 simulations, and colors indicate the frequency of SNVs within SAMs (darker the higher). Patterns whose average is more than 1.0 for S1 and S2, and 0.25 for F1 and F2 are shown.

### Patterns predicted by simulations using MAP parameter estimates

The predictive performance of the inferred models was further tested by stochastic simulations based on the MAP estimates of parameters (Table 1). This allowed us to evaluate various somatic SNVs patterns distinct from the inter-branch SNVs used for model fitting (Figs. 5, S12, S13).

Our simulations revealed that the topological congruence between somatic phylogenies and physical tree architectures is a key discriminator between the models of stem cell dynamics (Table 2). While conserved models predicted low congruence rates (0.144–0.480), replaced models yielded much higher congruence rates (0.228–0.876), closely matching the empirical observations where somatic SNVs largely reflect physical architectures (Satake *et al*., 2024). This high congruence in replaced models arises because somatic drift during apical growth aligns the genetic structure of the SAM with the physical branching history. This architectural alignment was further confirmed by unweighted Robinson-Foulds distances (Robinson & Foulds, 1981; Fig. S12). The distances were substantially smaller in replaced models, whereas conserved models generated large discordance not observed in the empirical data. Notably, the models also correctly predicted that, in smaller trees with fewer mutations (e.g., S2 and F2), topological deviations can frequently occur even under moderate drift, as observed in the F2 individual (Satake *et al*., 2024). This demonstrates that while somatic phylogenies reflect branching history, they are not strictly congruent with physical architecture, particularly when the number of accumulated mutations is insufficient to resolve the topology. These topological differences among models are partly driven by how somatic SNVs are distributed across branches (Fig. 5b).

**Table 2.** Topological congruence between phylogenetic and physical trees.

| Individual | Model I | Model II | Model III | Model IV |
| --- | --- | --- | --- | --- |
| S1 | 0.480 (120/250) | 0.440 (110/250) | 0.704 (176/250) | 0.876 (219/250) |
| S2 | 0.312 (78/250) | 0.284 (71/250) | 0.348 (87/250) | 0.548 (137/250) |
| F1 | 0.216 (54/250) | 0.220 (55/250) | 0.420 (105/250) | 0.456 (114/250) |
| F2 | 0.232 (58/250) | 0.144 (36/250) | 0.228 (57/250) | 0.348 (87/250) |

\* Congruence rate (out of 250 stochastic simulations).

The spatial distribution of somatic SNVs across branches further supports the replaced models (Fig. 5b). While the replaced models primarily predicted nested SNV distribution patterns that align with the tree topology, the conserved models also generated non-nested, “patchy” patterns (highlighted by asterisks in Fig. 5b). These patchy distributions were most prevalent in the conserved random model and less frequent in the conserved biased model, but were very few in the replaced models. Notably, our results show that biased branching produced fewer non-nested patterns than random branching. This effect of branching was not captured in Tomimoto & Satake (2023), which assumed that only lateral branches undergo sampling process by branching, whereas our current model incorporated branching at both lateral and main branches. Since these non-nested patterns were rarely observed in the Dipterocarpaceae trees (indicated by red asterisks in Fig. 5b), these distribution patterns provide independent support for the replaced models. Given that our parameters were estimated using distinct measurement (inter-branch SNVs), the ability of the replaced models to predict these spatial distributions highlights their robust predictive power.

We note, however, some discrepancies common to all models, which offer important insights into developmental constraints and sampling limitations. First, all models underestimated the number of somatic SNVs shared by all branches in individuals S2 and F1 (Fig. 5b). This underestimation suggests the significant number of mutation accumulation in the main trunk prior to the first bifurcation. Since our estimation framework is based on genetic differences between branch tips, such ancestral mutations shared across all branches are not fully captured in the current rate estimation. Indeed, all models also underestimated the somatic SNVs accumulated within the SAMs of all branches in individuals S2 and F1 (Fig. S13). Second, all models predicted a higher number of singletons than were empirically observed. This discrepancy likely reflects a detection limit in the genomic data. Namely, most singletons persist at low frequencies within the SAMs, as model predicted (Fig. 5b), and are not transmitted to sampled leaves, making them undetected by sequencing.

Our models provided a window into such internal genetic structures of the SAM that are not evident from current empirical data derived from leaf samples (Satake et al., 2024). All models predicted that approximately half of the accumulated SNVs reached fixation within the SAM, while the remaining half persisted at low frequencies, generating substantial genetic heterogeneity among stem cell lineages (Fig. S13). Again, on the total number of somatic SNVs, the replaced models produced better predictions for individuals S1 and F2, whereas the conserved models underestimated the accumulation in all individuals. While physical architecture alone could not account for the elevated mutation load in individuals S2 and F1, these results suggest that somatic SNVs can serve as a molecular record of hidden growth histories, capturing periods of increased mutation accumulation during early development that leave no discernible morphological trace.

## DISCUSSION

In this study, we successfully translated snapshot genomic variation into the dynamic processes of mutation accumulation during growth of long-lived Dipterocarpaceae trees, which dominate tropical rain forests in Southeast Asia (Ghazoul, 2016). We focused on somatic genetic drift arising from the replacement of stem cell lineages within the shoot apical meristem (SAM) during both apical growth and branching (Fig. 1). To quantify this process, we analyzed inter-branch somatic SNVs—genetic differences between branch tips—in relation to the physical distance separating branches along the branching architecture. This relationship can be summarized by a linear like function, whose intercept reflects the degree of somatic genetic drift (Fig. 3). Under strong somatic drift, mutations rapidly fix within the SAM and spread to both branches, such that only mutations arising after a forking event contribute to genetic differences between branches, resulting in an intercept close to zero. In contrast, under weak somatic drift, genetic heterogeneity is maintained within the SAM, and mutations accumulated before forking also contribute to genetic differences between branches, producing a positive intercept.

The strength of somatic drift is determined by the dynamic processes of apical growth and branching, as well as by key parameters such as the number of stem cells (𝛼) and the rate of cell division per unit length of growth (𝑟). In addition, the mutation rate per unit length (𝑢) determines the slope of the relationship (Fig. 3b). By fitting this model to the observed data, we inferred somatic drift and mutation rates in the Dipterocarpaceae trees.

### Moderate somatic genetic drift during apical growth of Dipterocarpaceae trees

Model fitting using ABC-SMC inferred a moderate degree of somatic drift in the tropical trees. Both the fitting results and the somatic SNVs patterns predicted by stochastic simulations were most consistent with the models of replaced apical growth (Figs. 3, 5, S6–9A, S10, S13; Table 2). These results indicate that both apical growth and branching contribute to somatic drift in the Dipterocarpaceae trees. Although random branching was favored by model fitting (Figs. S8A, S9A, S14A), the predicted patterns were very similar between branching models, preventing us from excluding biased branching. This may reflect the dominant role of apical growth in shaping somatic drift in these tropical trees. While stem cell lineages in the SAM are largely conserved in the annual plant *Arabidopsis thaliana* (Burian *et al*., 2016), developmental traces of somatic mutations in Dipterocarpaceae trees suggest frequent lineage replacement over decades of apical growth.

We note, however, that our study assumes branching occurs exclusively at bifurcation points, with apical growth occurring along branches between them. This assumption may overlook branching events that occur along elongating shoots. In such cases, somatic drift attributed to apical growth could partially result from branching processes. In natural trees, however, detecting such events is difficult because their developmental traces may disappear over long growth periods (Barthélémy & Caraglio, 2007).

Additionally, the detailed parameter values governing somatic drift, such as 𝛼 and 𝑟, remain uncertain in our approach. Accordingly, we emphasize the inferred degree of ‘moderate’ somatic drift rather than precise parameter estimates. As indicated by the MSE landscapes, 𝛼 and 𝑟 are correlated in replaced models such that 𝛼/𝑟 ≈ const, reflecting a certain degree of somatic drift (Figs. S8–9D). Empirical estimation of these parameters could enable more precise inference (Watson *et al*., 2016; Yu *et al*., 2024), although such measurements remain challenging in long-lived trees. For instance, Yu *et al*., (2024) estimated the number of stem cells in the L1 layer of eelgrass to be 7–12 cells, improving their model predictions. If 𝛼 = 10 in Dipterocarpaceae trees, our ABC-SMC analysis assigns the highest posterior probability to the replaced random model (Fig. S14), with a MAP estimate of the cell division rate of 𝑟 ≈ 4. This rate is an order of magnitude lower than that estimated for *A. thaliana* (Watson *et al*., 2016), potentially reflecting evolutionary mechanisms that suppress mutational load in long-lived plant (Klekowski, 2003; Lanfear *et al*., 2013).

In contrast to our inference of moderate somatic drift in Dipterocarpaceae trees, where a large fraction of somatic SNVs were inferred to be fixed within a SAM (Fig. S12), Schmitt *et al*. (2024) reported abundant low-frequency SNVs within the SAMs of *Dicorynia guianensis* and *Sextonia rubra* in an Amazonian forest. These observations suggest very weak somatic drift during growth of these species. Such weak drift is also consistent with the observed topological incongruence between somatic phylogenies and physical architectures, as predicted by conserved models in our study (Tabel 2; Fig. S12; Tomimoto & Satake, 2024). Together, these findings indicate that strength of somatic drift may vary substantially among plant species, highlighting the importance of species-specific inference of stem cell dynamics.

We note, however, that differences between the two studies may partly arise from differences in sampled tissues, i.e., leaves in Satake *et al*. (2024) versus buds in Schmitt *et al*. (2024). While a bud contains stem cells from all meristematic layers, a leaf originates from a subset of sectorial stem cells (Irish & Sussex, 1992; Frank & Chitwood, 2016) and predominantly consists of L2 derived cells (Dermen, 1953; Poethig, 1989), which may reduce within-sample genetic variation in Satake *et al*. (2024). Consequently, our model may primarily capture stem cell dynamics within the L2 layer, whereas the dynamics across layered structure remained unresolved. Notably, using layer-specific samples, Goel *et al*. (2024) demonstrated layer-specific accumulation of somatic mutations in an apricot tree and observed somatic phylogenies that perfectly reflected physical architecture within each layer. Their results suggest frequent lineage replacement within layers but not across layers. If similar processes also occur in both Satake *et al*. (2024) and Schmitt *et al*. (2024), differences in topological congruence may largely reflect difference in tissue sampling.

### Mutation rates and genetic structures shaped by somatic drifts

By accounting for stem cell dynamics, our model inferred somatic mutation rates in Dipterocarpaceae trees (Fig. 3a). Across all models, inferred mutation rates were lower than previous estimates, suggesting that approaches that ignore stem cell dynamics may overestimate mutation rates (Satake *et al*., 2024). However, the conserved models, which inferred particularly low rates, may underestimate mutation rates in this system, as they predicted unrealistically low numbers of accumulated SNVs within branches (Fig. S13). In contrast, replaced models provided more accurate predictions of SNV accumulation per branch, together with consistent other somatic SNV patterns and superior model fit, as discussed above. These results suggest that replaced models yield more reliable mutation rate estimates (Fig. 4a; Table 1).

Nevertheless, even replaced models predicted lower SNV accumulation in branches of individuals S2 and F1 (Fig. S12). This discrepancy may arise because a substantial fraction of mutations accumulated before the first branch bifurcated from the trunk, as suggested by deviations in somatic SNV distribution patterns (Fig. 5b). Dipterocarpaceae seedlings are shade tolerant and can persist as suppressed seedlings for extended periods, sometimes for decades, while awaiting canopy gap formation (Scholes *et al*., 1996). During this suppressed phase, mutations may accumulate without corresponding increases in tree height or trunk diameter. Individuals S2 and F1 may thus have underwent prolonged suppressed growth, leading to elevated early mutation accumulation without leaving a clear developmental trace in their physical appearance. As tropical trees do not produce annual rings, estimating this suppressed-growth phase is technically difficult, but somatic SNVs provide insights into such developmental events.

Somatic mutations allow us to trace the developmental history of long-lived plants, particularly the behavior of stem cells within the SAM (Poethig, 1989; Ji *et al*., 2025). Prior to reproduction, SAMs transition into inflorescence meristems, and stem cells in these meristems give rise to gametes such as pollen and ovules (Ji *et al*., 2025). Thus, somatic mutations accumulated in SAMs during growth can be transmitted to offspring (Plomion *et al*., 2018; Wang *et al*., 2019; Schmitt *et al*., 2024; Goel *et al*., 2024), directly contributing plant evolutions (Whitham & Slobodchikoff, 1981; Schoen & Schultz, 2019; Smith *et al*., 2026). From an evolutionary perspective, stem cells in SAMs thus correspond to germline cell lineages in mammals (Frumkin *et al*., 2005; Reizel *et al*., 2012). Our results revealed how the lineages are maintained during the hundreds of years development of Dipterocarpaceae trees. Namely, multiple stem cell lineages within a SAM are preserved only in the short term due to lineage replacement by somatic drift, coalescing to a single lineage during apical growth. In the long term, lineages are kept separate by branching architecture. Germline lineages that independently accumulate mutations are thus mainly separated by physically distinct branches, while relatively shorter lineages are separated within a SAM, producing genetic heterogeneity among one-half of stem cells (Fig. S13).

Stochastic replacement of stem cell lineages within a SAM may facilitate somatic selection, or within-individual selection, among cells in Dipterocarpaceae trees.

Previous modeling studies have shown that somatic selection can accelerate the fixation of advantageous mutations within a SAM (Pineda-Krch & Fagerström, 1999), while purging deleterious mutations (Klekowski & Kazarinova-Fukshansky, 1984b; Otto & Orive, 1995), potentially supporting the longevity (Ally *et al*., 2010). In addition, selection among modules can enhance individual-level adaptation to herbivores and pests

(Antolin & Strobeck, 1985; Gill *et al*., 1995; Orive, 2001; Folse & Roughgarden, 2012). Despite these theoretical predictions, Satake et al. (2024) and most empirical studies have found no evidence for somatic selection within trees (Orr *et al*., 2020; Perez-Roman *et al*., 2022; Duan *et al*., 2022), motivating our assumption of neutrality for SNVs. Nonetheless, the support for replaced apical growth in our analysis raises the possibility that somatic selection operates within SAMs. Because detecting such selection is statistically challenging, future progress will require further integration of empirical data with mathematical and computational modeling approaches (Reusch *et al*., 2021).

### Conclusions and future perspective

Our model successfully inferred the dynamics of somatic mutation accumulation in Dipterocarpaceae trees, revealing a moderate degree of somatic genetic drift during apical growth and mutation rates slightly lower than previous estimates that ignored stem cell dynamics. Using the estimated parameters, we conducted stochastic simulations that predicted patterns of somatic SNVs not directly observed in the sampled trees. In particular, somatic phylogenies often deviated from physical architecture, especially in smaller individuals. The simulations further indicated that approximately half of the accumulated SNVs became fixed within a SAM, while the remaining SNVs persisted at low frequency, generating genetic heterogeneity among stem cells.

Beyond explaining observed patterns, our framework enables the prediction of somatic SNV patterns in non-sequenced branches within individuals, given their branching architecture. This approach can be extended to all individuals of the same Dipterocarpaceae trees within the forest, provided that their age and branching architecture are known. Thus, even without sequencing every individual, our model offers a way to evaluate the amount of standing genetic variation arising from somatic mutations at the forest scale by integrating branching architecture and demographic history.

Nevertheless, our model has limitations in estimating detailed parameters governing somatic drift. Under replaced apical growth, the number of stem cells, 𝛼, and the rate of cell division, 𝑟, are not separately identifiable. At least one of these parameters must therefore be constrained empirically by direct histological observation (Yu *et al*., 2024), although this is feasible only for a limited number of species and often difficult to obtain for long-lived trees. Alternatively, the site frequency spectrum of somatic mutations within a SAM could be used to infer 𝛼 (Johannes, 2025). Although our framework allows to directly derive the spectrum from the mutational state vector (SI Methods S1), this approach is not applicable to leaf-based sequencing data from the Dipterocarpaceae trees. This limitation also underscores that the choice of summary statistics for model fitting remains a critical consideration, depending on the available tissue types and sequencing depth. Finally, Satake et al. (2024) suggested that somatic mutations accumulate in a time-dependent rather than cell division–dependent manner. Testing this hypothesis while explicitly accounting for stem cell dynamics will require independent estimation of cell division rates, which remains an important challenge for future studies.

To fully understand the diversity of somatic mutation dynamics across plant systems, it will be essential to apply this framework to a broader range of empirical datasets spanning different species, life histories, and ecological conditions. Such comparative analyses will provide a general understanding of how developmental processes, growth environments, and life-history strategies shape the accumulation of somatic mutations in long-lived organisms. Ultimately, integrating empirical genomic data with mechanistic modelling frameworks offers a powerful approach to uncover universal principles governing somatic evolution in plants.

## Acknowledgements

We thank the following people for their helpful comments: R. Hayashi, R. Imai, Y. Iwasa, S.N. Kudo, K. Matsuo, K. Noshita., S.P. Otto, E. Sasaki., and S. Yeaman. We also thank ChatGPT (OpenAI) and Gemini (Google) for their assistance in English language editing and for providing constructive feedback on the structure of the manuscript. This work was supported by JSPS KAKENHI Grant Number 23KJ1722 to S.T., and Grant Number 23H04966 to A.S.

## Author Contributions

**S. T.** Conceptualization, Methodology, Visualization, Writing – original draft, Writing – review & editing, Funding acquisition; **A.S.** Conceptualization, Writing – review & editing, Supervision, Funding acquisition.

## Data Availability Statement

The data used in this research are based on the published paper (Satake *et al*., 2024). All Python codes would be available from the corresponding author upon reasonable request.

## Online Supporting Information

Additional supporting information may be found in the Supporting Information section at the end of the article.

## Supporting Information for “Inferring somatic mutation dynamics from within-individual genomic variation across branches in tropical trees” by Sou Tomimoto and Akiko Satake

### Methods S1

#### Modelling somatic mutation accumulation by Markov chain

During growth, a tree expands its branches and accumulates somatic mutations, leading to genetic difference among branches within an individual. The accumulating pattern of somatic mutations is closely linked to the growth processes (i.e., somatic genetic drift) of a tree. Here, we introduce a mathematical model that simulate the dynamic process of somatic mutation accumulation in a tree with a given branching architecture, modifying previous work by (Tomimoto & Satake, 2023).

The model focuses on mutations accumulating in the stem cells of a shoot apical meristem (SAM). As the stem cells originate all aerial parts of a tree (Stewart & Dermen, 1970), mutations accumulated in these stem cells may determine the genetic structure within a tree (Schmid-Siegert *et al*., 2017). Stem cells in a SAM undergo asymmetric cell divisions, producing a successor stem cell and a non-successor cell that enters the differentiation pathway. These non-successor cells contribute to the formation of a phytomere—a growth unit consisting of a node with its attached leaf, an internode below, and an axillary bud at the internode’s base (White, 1979; Evert & Eichhorn, 2006). Branch elongates through the repetitive production of phytomeres, we call this process as “apical growth,” alongside the proliferation and expansion of internode cells (Burian et al., 2016; McKim, 2020).

#### Biological assumptions in the model

The model focuses on mutations accumulating in stem cells of a SAM. As the stem cells originate all areal part of a tree, mutations accumulated in these stem cells may determine the genetic structure within a tree (Schmid-Siegert *et al*., 2017). Stem cells in a SAM undergo asymmetric cell divisions, producing a successor stem cell and a non-successor cell that enters the differentiation pathway. These non-successor cells contribute to the formation of a phytomere—a growth unit consisting of a node with its attached leaf, an internode below, and an axillary bud at the internode’s base (White, 1979; Evert & Eichhorn, 2006). Branch elongation, or apical growth, occurs through the repetitive production of phytomeres, alongside the proliferation and expansion of internode cells (Godin & Caraglio, 1998; Evert & Eichhorn, 2006; Burian *et al*., 2016). During this ‘apical growth’ process, called ‘elongation’ in the previous study (Tomimoto & Satake, 2023), mutations may accumulate in successor stem cell lineages. If a stem cell lineage fails to leave a successor stem cell, other stem cell lineage replaces the lineage, and mutations accumulated in the lineage are lost from a SAM. Such replacement of stem cell lineage leads to somatic genetic drift (Schoen & Schultz, 2019), and the strength of somatic drift during apical growth may differ between species (Stewart & Dermen, 1970; Ruth *et al*., 1985; Rogers & Bonnett, 1989; Burian *et al*., 2016). In the present study, the conserved and replaced lineage model describes the absence and presence of somatic drift in apical growth (Fig. 1a). Note that the previous study called each model as called structured and stochastic model, respectively (Tomimoto & Satake, 2023). In the conserved model, 𝛼 stem cells in a SAM produce 2𝛼 daughter cells through cell division. One daughter cell from each stem cell remains in the SAM while the other differentiates. Thus, asymmetric cell divisions occur successfully, conserving all stem cell lineages during apical growth. In the replaced model, subsequent stem cells are randomly sampled from 2𝛼 daughter cells, resulting in the stochastically replacement of lineages.

Along with apical growth, a SAM produces new meristems at the base of internodes. These axillary meristems can grow into lateral branches, although most of them remain as dormant buds (Poethig, 1989; Cline, 1997; Sussex & Kerk, 2001). In this process, called ‘branching,’ some of the stem cell lineages contribute to the axillary meristem formation, and mutations accumulated in these lineages are passed on to the new meristem (Burian *et al*., 2016). Branching is analogous to sampling mutations accumulated in the main branch for a new branch. The way of sampling may differ among species depending on the developmental mechanism in the axillary meristem formation, resulting in different strength of bottleneck effect, i.e., somatic genetic drift (Yu *et al*., 2020). In the present study, the ‘biased’ and ‘random’ sampling describes the weak and strong somatic genetic drift in branching, respectively (Fig. 1b). In random branching, stem cells in the apical meristem contribute to the formation of the axillary meristem with equal probability, allowing sampling of multiple cell lineages. Contrastingly, in biased branching, only a subset of stem cells in the apical meristem contribute to the formation of the axillary meristem (Irish & Sussex, 1992; Burian *et al*., 2016). A single cell lineage is sampled to for a new branch, resulting in strong bottleneck effect in branching.

#### Modelling a distribution of the number of mutated cells in a SAM

We first model mutational state within a SAM based on the Markov chain framework (Klekowski *et al*., 1989; Tomimoto & Satake, 2023). The model focuses on the number of mutated cells at a focal site within a SAM. The probability of a SAM having *i* mutated cells at time *t* is denoted by 𝜋_1_(𝑡). For a SAM consisting of 𝛼 stem cells, 𝛼 + 1 possible states are considered, and each state is given by an element of a state vector as follows:

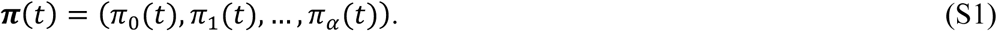

The first element of the vector 𝜋_0_(𝑡), for instance, represents the probability of a SAM having no mutated cell, and the last 𝛼 + 1th element 𝜋_α_(𝑡) represents the probability of a SAM consisting only of mutated cells (i.e., mutated cells are fixed within the stem cell population). The state vector represents the probability distribution of the number of mutated stem cells within a SAM. For simplicity, we assumed that the number of stem cells remains constant throughout the growth (i.e., the number of elements in 𝝅(𝑡) remains constant).

At the beginning of a tree development, no somatic mutations accumulate in a SAM with probability one, 𝜋_0_(𝑡) = 1. The initial condition of the state vector is given as 𝝅(0) = (1, 0,…, 0). From this initial state, somatic mutations occur and accumulate in a SAM during growth process. Assuming that somatic mutations occur independently in each growth process, the inhomogeneous Markov chain describes the change in mutational state in a SAM during apical growth and branching by transition matrices 𝑳 and 𝑩, respectively. For instance, the state of a SAM that underwent 𝑡_0_times cell divisions by apical growth from the seedling is given as 𝝅(𝟎)𝑳t_0_. If the SAM underwent additional branching and 𝑡_1_ times cell divisions of apical growth, the state is now given by 𝝅(𝟎)𝑳t_0_𝑩𝑳t_1_. This gives the mutational state of br-1 branch in Fig. 1c. We modelled transition matrices 𝑳 for apical growth with and without stem lineage replacement and 𝑩 for branching with random and biased sampling, extending formulas in the previous studies (Klekowski *et al*., 1989; Tomimoto & Satake, 2023).

### Transition matrix of apical growth

To express the processes of apical growth that differs in the degree of somatic genetic drift, two contrasting transition matrices are introduced based on the previous studies (Klekowski *et al*., 1989; Tomimoto & Satake, 2023). In the first ‘conserved’ model, stem cell lineages are conserved during the process of apical growth without the drift (Fig. 1a), which was called “structured elongation” in the previous studies. In the second “replaced” model, the replacement of stem cell lineages randomly occurs during apical growth with the strong drift (Fig. 1b), which was called ‘stochastic elongation’ in the previous studies. The conserved model maintains all stem cell lineages within a SAM. During cell division, 𝛼 stem cells produce 2𝛼 daughter cells. One of the daughter cells from each stem cell persists in a SAM, while the other daughter cells differentiate and leave the SAM (i.e., asymmetric cell divisions occur successfully). Mutation occurs during cell divisions and accumulate in each lineage. Thus, changes in the number of mutated stem cells occur solely due to mutations. The transition probability for the number of mutated cells changing from 𝑖 to 𝑗 is given as follows:

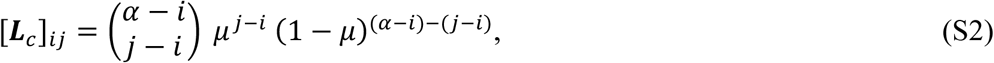

where 𝜇 is the mutation rate per cell division per site. Here, [𝑨]*_ij_* represents the 𝑖𝑗 element of matrix 𝑨. Eq. (S2) gives the probability that 𝑗 − 𝑖 cells mutates while others remain non-mutated. Eq. (S2) holds for 𝑗 ≥ 𝑖; otherwise, [𝑳_c_]_ij_ = 0.

The replaced model permits the replacement of stem cell lineages. Cell division produces 2𝛼 daughter cells, from which 𝛼 cells are randomly sampled for the next stem cells, leading to the random replacement of stem cell lineages. Consequently, changes in the number of mutated stem cells occur due to both mutations and lineage replacement. The transition probability for the number of mutated cells changing from 𝑖 to 𝑗 is given as follows:

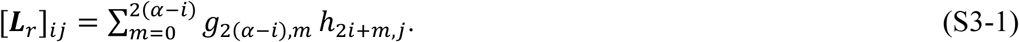

Here, 𝑔_2(a-i),m_ is the probability that 𝛼 − 𝑖 non-mutated cells produce *m* mutated cells thorough single cell division and given as follows:

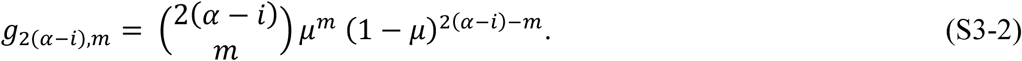

This equation shows the probability that, of 2(𝛼 − 𝑖) daughter cells produced from the non-mutated cells, 𝑚 daughter cells mutate. Given 𝑚 daughter cells newly mutated, the number of mutated daughter cells becomes 2𝑖 + 𝑚 after a cell division. The probability that 𝑗 mutated cells are sampled from 2𝑖 + 𝑚 daughter cells and 𝛼 − 𝑗 non-mutated cells are sampled from 2𝛼 − (2𝑖 − 𝑚) daughter cells is given as

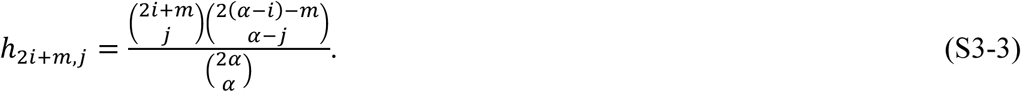

Together, the probability that the number of mutated cells changes from 𝑖 to 𝑗 is given as the product for all possible case as given in Eq. (S3-1). Refer to Klekowski *et al*. (1989).

### Transition matrix of branching

In addition to the apical growth process, two transition matrices for branching are introduced that differ in the degree of somatic genetic drift. In the first “random (unbiased)” model, stem cells for a new axillary meristem are randomly sampled from the apical meristem. This allows the multiple stem cell lineages to participate in the formation of new axillary meristem, leading to a weak somatic drift. In the second “biased” model, a new axillary meristem is formed from a single stem cell lineage in the apical meristem, leading to stronger somatic drift. To focus on the somatic drift in branching, the occurrence of *de novo* mutations during branching is ignored here (but see Burian *et al*. (2016) and Lanfear (2018) for the discussions on un-negligible cases).

In the random branching, the transition probability of the number of mutated cells from 𝑖 in the apical meristem to 𝑗 in axillary meristem is given as follows:

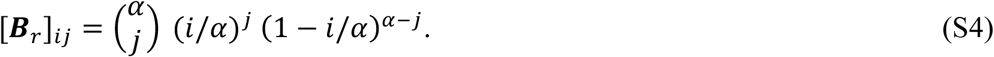

Here, 𝑖/𝛼 and 1 − 𝑖/𝛼 are the fraction of mutated and nonmutated stem cells in the apical meristem, respectively. Sampling probability depends on these fractions. Note that Eq. (S4) implies the sampling of stem cells from a large number of cells, ignoring the spatial effect which result in biased sampling (Tomimoto & Satake, 2023).

To describe the biased sampling, we introduced a transition matrix as follows:

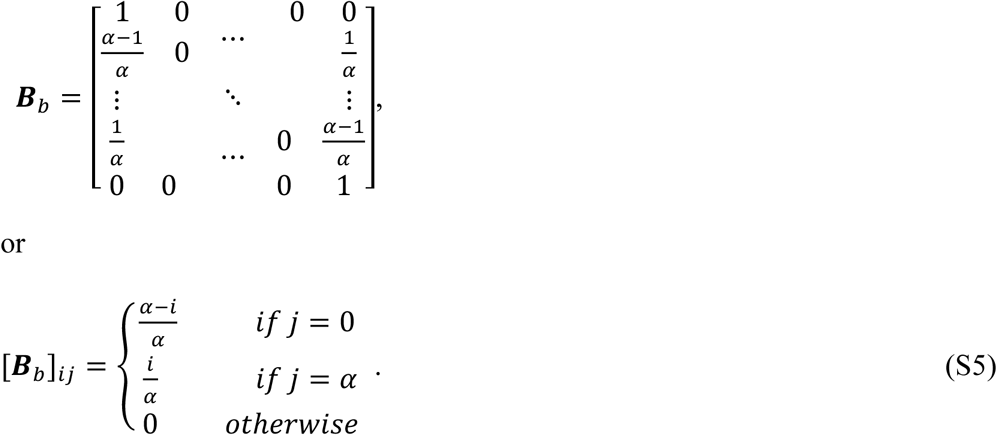

In this biased branching, the transition occurs only to state 0 or 𝛼. After branching, the mutant cells are either lost or fixed in the axillary meristem, and the probabilities depends on the fraction of the mutant cells in the apical meristem. Thus, Eq. (S5) assumes that an axillary meristem is formed from a single-stem cell lineage in the apical meristem, i.e., biased sampling.

### Methods S2

#### Inter-branch SNVs: genetic difference between two branches

We define the genetic difference by heterozygosity between cells that are randomly sampled from the SAM of each branch. Given there are 𝑖 and 𝑗 mutated cells at branches 𝑛 and 𝑚, respectively, the heterozygosity is given by:

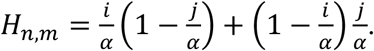

Here, the first term 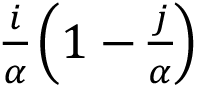 is the probability of sampling a mutated cell from branch 𝑛 and sampling a non-mutated cell from branch 𝑚, while the second term corresponds to the opposite case. Then, the probability of inter-branch genetic difference at a focal site is:

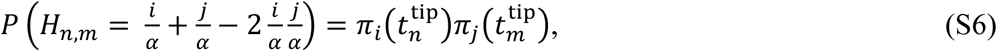

where 𝜋_i_(𝑡_n_^ip^) is the probability that the tip of branch 𝑛 harbors 𝑖-th mutated cells. Thus, the average heterozygosity is given by:

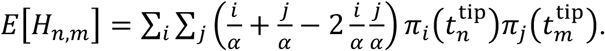

Because the two branches share their developmental history before the forking point, their mutational states at the tips, 𝜋_i_(𝑡_n_^tip^)and 𝜋_j_(𝑡_m_^tip^), are generally correlated. However, the two branches evolve independently and follows the Markov process, conditional on the mutational state at the forking point 𝑡*_n,m_^fork^*. To apply our Markov chain framework, we conditionalize the average heterozygosity by the mutational state at forking point 𝑆(𝑡*_n,m_^fork^*), where 𝑆(𝑡) represents the number of mutated stem cells at growth point 𝑡 and its probability is given by 𝑃(𝑆(𝑡) = 𝑙) = 𝜋_L_(𝑡). Namely,

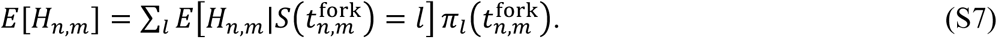

The conditional average heterozygosity is given by:

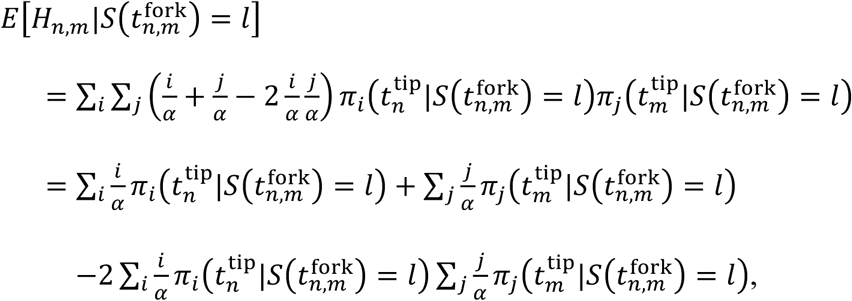

where, in the last line, the independence of branches after forking is used. Therefore, conditional average heterozygosity is simplified as follows:

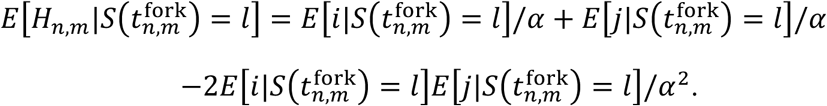

Thus, plugging this into Eq. (S7), we get:

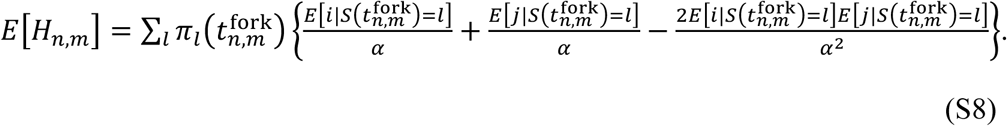

Since 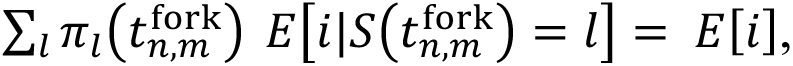 the equation is simplified into:

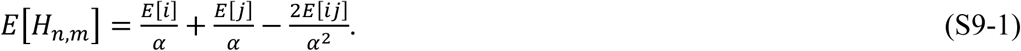

where,

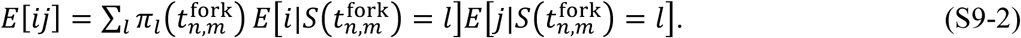

Here, we note that the expectations are derived from the state vectors of the Markov chain model: 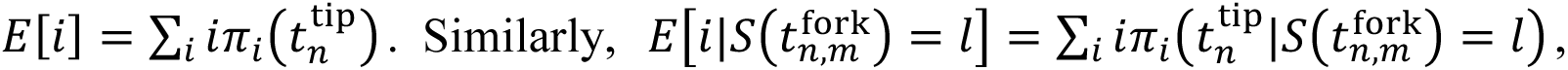 where 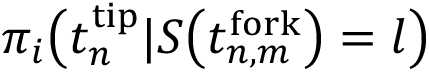 is the 𝑖-th element of the state vector initialized under the condition 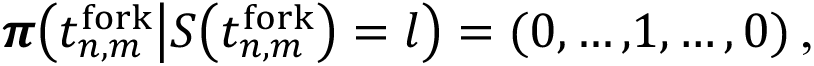 whose 𝑖-th element is given by 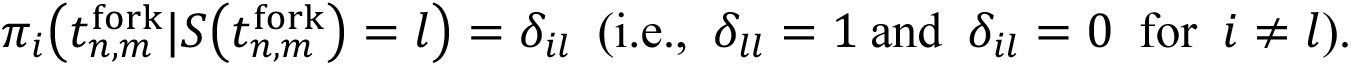

Taking Fig. 1c as an example, consider the genetic differences between branch br-1 and br-2. The state of br-1 branch initialized under the no mutations condition at forking point is given by: 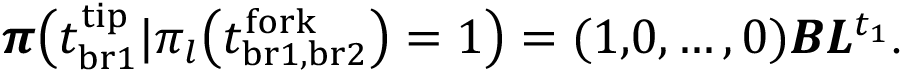 For all *i*-th state conditions, we can get the state of br-1 branch, by changing the initial state. Similarly, we can get conditional state of br-2 branch. Additionally, 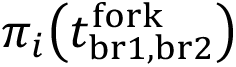 is given by the probability of state 𝑖 at the forking point, i.e., 𝑖-th element of 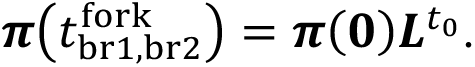 In this way, the model prediction of inter-branch SNVs is calculated from by Eq. (S8).

---

**Table S1.** Branching architectures.

| Symbol | Values for S1 | Values for S2 | Values for F1 | Values for F2 |
| --- | --- | --- | --- | --- |
| s0 | 21.79 | 22.2 | 23.71 | 25.27 |
| s1 | 8 | 3.3 | 4.4 | 2.1 |
| s2 | 10 | 3.6 | 6.4 | 3.8 |
| s3 | 11.75 | 7.5 | 13.1 | 8.9 |
| b10 | 15 | 4.1 | 6.35 | 1.1 |
| b11 | 14.18 | 9.25 | 9.65 | 8.2 |
| b12 | 4.74 | 9.1 | 4.8 | 5.8 |
| b20 | 11.3 | 6.8 | 5.4 | 10.4 |
| b21 | 8.9 | 3.6 | 3.1 | 2.4 |
| b22 | 8.1 | 3.3 | 2.75 | 2.4 |
| b30 | 2.05 | 5.35 | 6.05 | 6.45 |
| b31 | 3.2 | 2.7 | 1.5 | 2.8 |
| b32 | 2.9 | 2.4 | 1.4 | 3.3 |

\*Values are from Satake *et al.* (2024).

**Table S2.** Other parameters in the simulation models.

| Symbol | Definition | Values |
| --- | --- | --- |
| $r_b$ | Number of cell divisions during branching | 7 (from Burian et al., 2016) |
| $\sigma$ | Degree of sampling bias during branching | 0.5 (Biased) or<br>10 (Random) |
| $u$ | Center position of newly formed branch | $u \sim U(0, 2\pi)$ |
| $G_{S1}$ | Genome size for S1 | 388,801,756 |
| $G_{S2}$ | Genome size for S2 | 320,739,335 |
| $G_{F1}$ | Genome size for F1 | 323,729,573 |
| $G_{F2}$ | Genome size for F2 | 263,488,812 |
| $v_i$ | Mutation rate per whole genome per cell division for individual $i \in \{S1, S2, F1, F2\}$ | $\mu_i G_i$ (where $\mu_i = u_i/r$) |

## Supplementary figures

**Figure S1.**
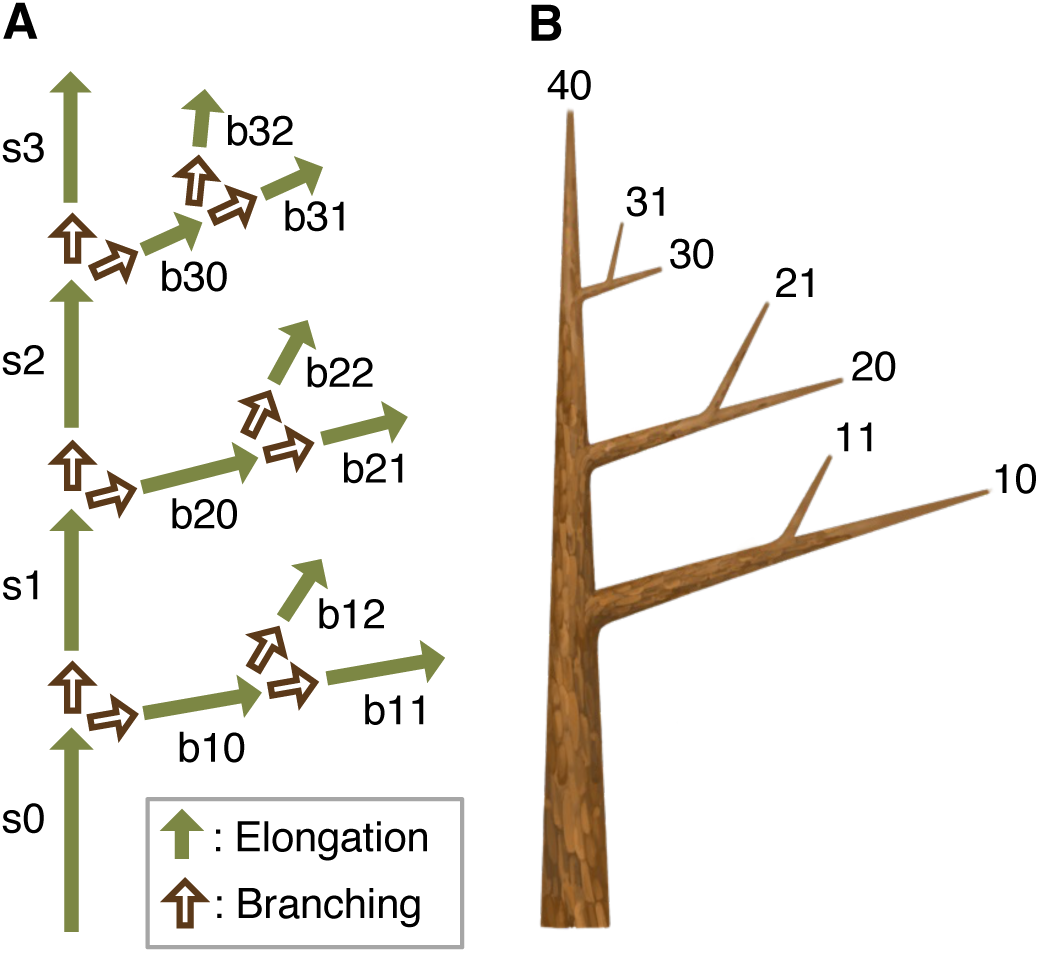
Branching architecture of trees. (A) Assumed growth processes of the tropical trees. The green-filled arrow represents the apical growth process. The symbols next to the arrow represent the length of the corresponding branch (i.e., for the branch of length s0, the SAM undergoes 𝑳^HTIIJ(5:!)^ apical growth, where floor(·) is the floor function). The values are listed in Table S1. The brown-unfilled arrow represents the branching process. Both main and lateral branches undergo the process of branching. (B) Corresponding branching architecture of a tree and ID of branch tips.

**Figure S2.**
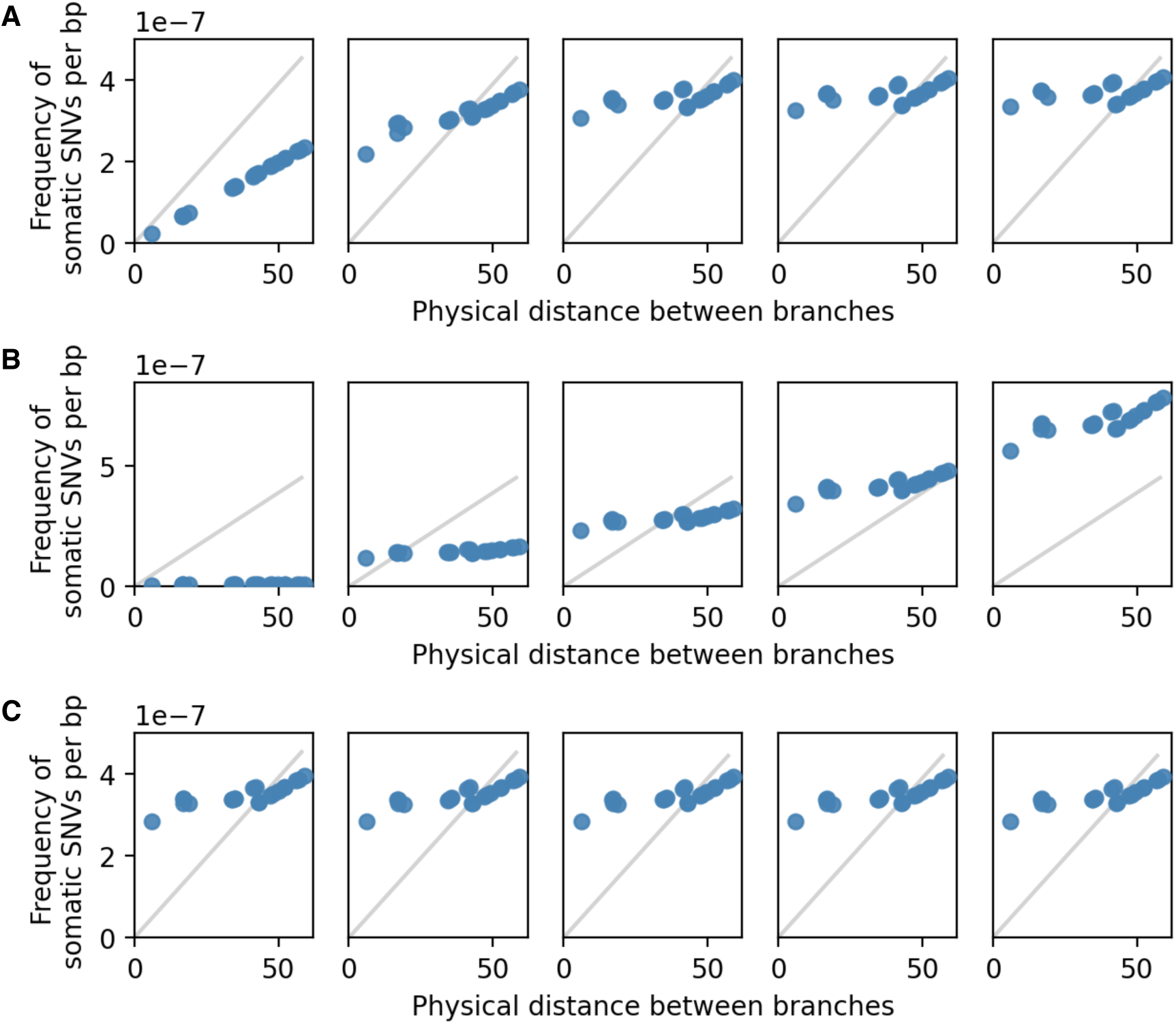
Parameter dependence of the model I (conserved random model). (A) Dependence on the number of stem cells. The values are 𝛼 = 1, 5, 15, 25, and 35, corresponding to the order of the subfigures from left to right. (B) Dependence on mutation rate per unit distance growth. The values are 𝑢 = 10^-10^, 1.7×10^-9^, 3.3×10^-9^, 4.9×10^-9^, and 8.0×10^-9^. (C) Dependence on the number of cell divisions per meter. The values are 𝑟 = 1, 3, 13, 23, and 33. While varying one parameter, other parameters were fixed with 𝛼 = 10, 𝑢 = 4.0×10^-9^, and 𝑟 = 5. The values highlighting parameter dependency were selected. The grey line is shown for comparison and comes from the regression of sequence data.

**Figure S3.**
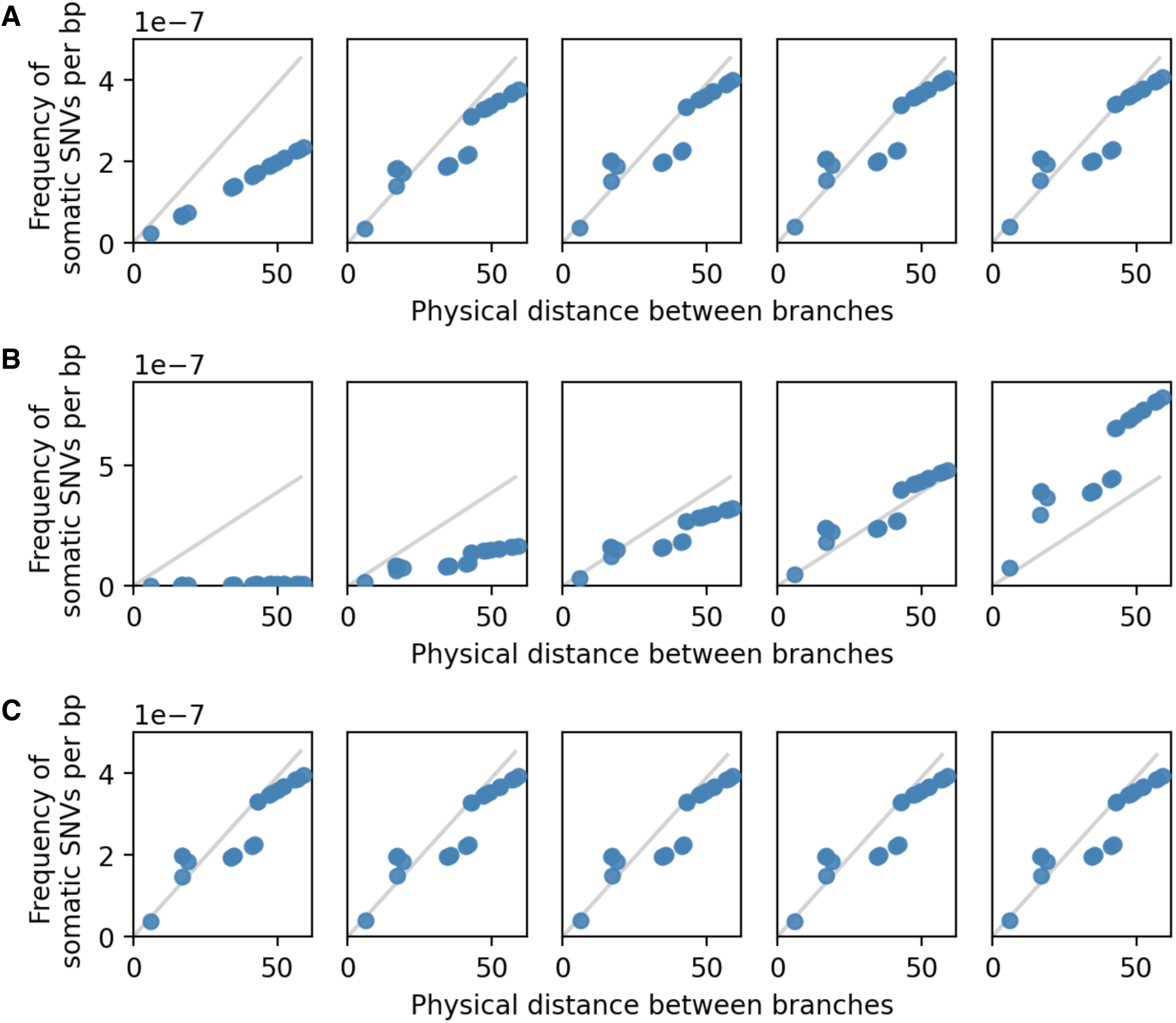
Parameter dependence of the model II (conserved biased model). (A) Dependence on the number of stem cells. The values are 𝛼 = 1, 5, 15, 25, and 35, corresponding to the order of the subfigures from left to right. (B) Dependence on mutation rate per unit distance growth. The values are 𝑢 = 10^-10^, 1.7×10^-9^, 3.3×10^-9^, 4.9×10^-9^, and 8.0×10^-9^. (C) Dependence on the number of cell divisions per meter. The values are 𝑟 = 1, 3, 13, 23, and 33. While varying one parameter, other parameters were fixed with 𝛼 = 10, 𝑢 = 4.0×10^-9^, and 𝑟 = 5. The values highlighting parameter dependency were selected. The grey line is shown for comparison and comes from the regression of sequence data.

**Figure S4.**
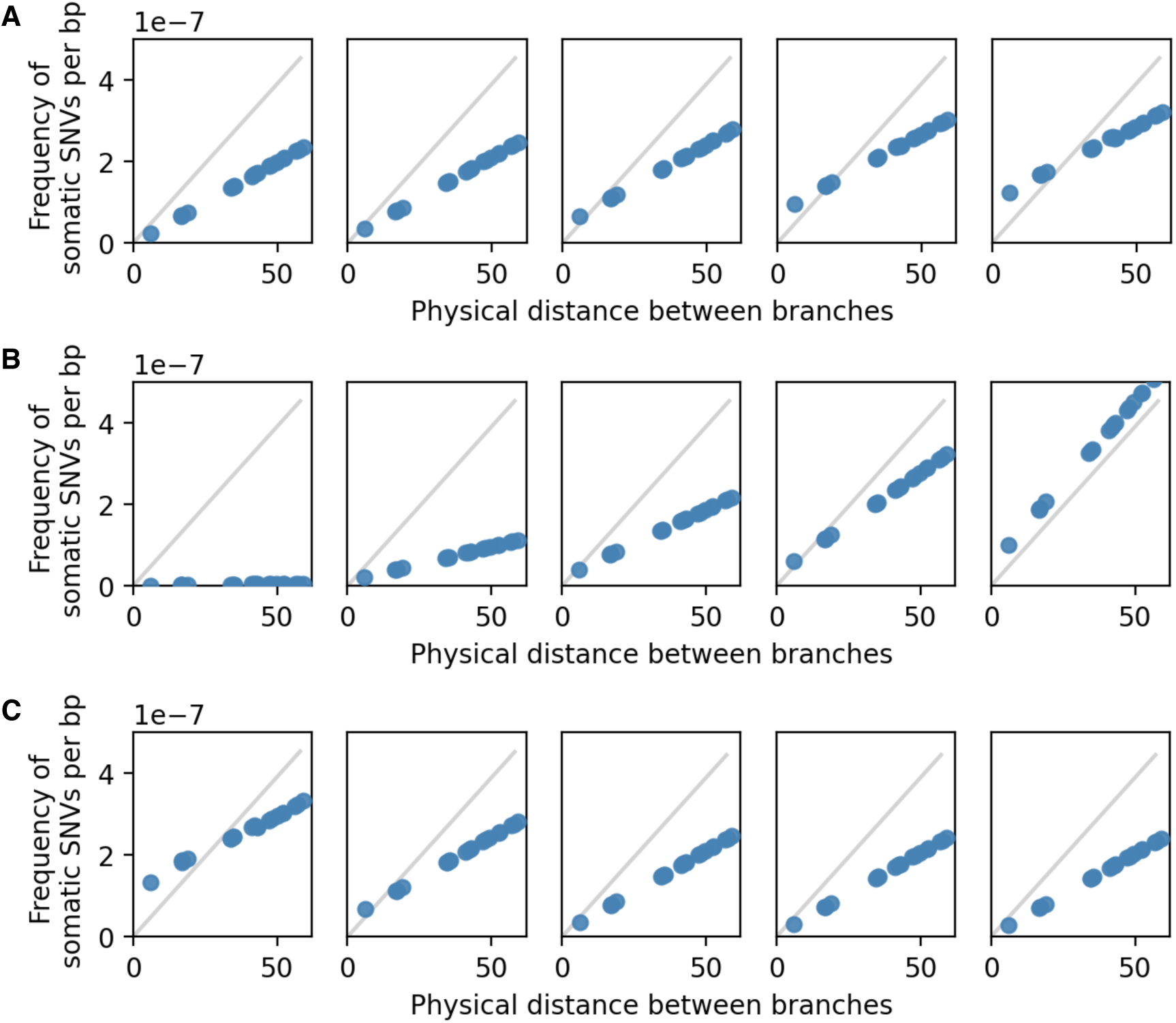
Parameter dependence of the model III (replaced random model). (A) Dependence on the number of stem cells. The values are 𝛼 = 1, 5, 15, 25, and 35, corresponding to the order of the subfigures from left to right. (B) Dependence on mutation rate per unit distance growth. The values are 𝑢 = 10^-10^, 1.7×10^-9^, 3.3×10^-9^, 4.9×10^-9^, and 8.0×10^-9^. (C) Dependence on the number of cell divisions per meter. The values are 𝑟 = 1, 3, 13, 23, and 33. While varying one parameter, other parameters were fixed with 𝛼 = 10, 𝑢 = 4.0×10^-9^, and 𝑟 = 5. The values highlighting parameter dependency were selected. The grey line is shown for comparison and comes from the regression of sequence data.

**Figure S5.**
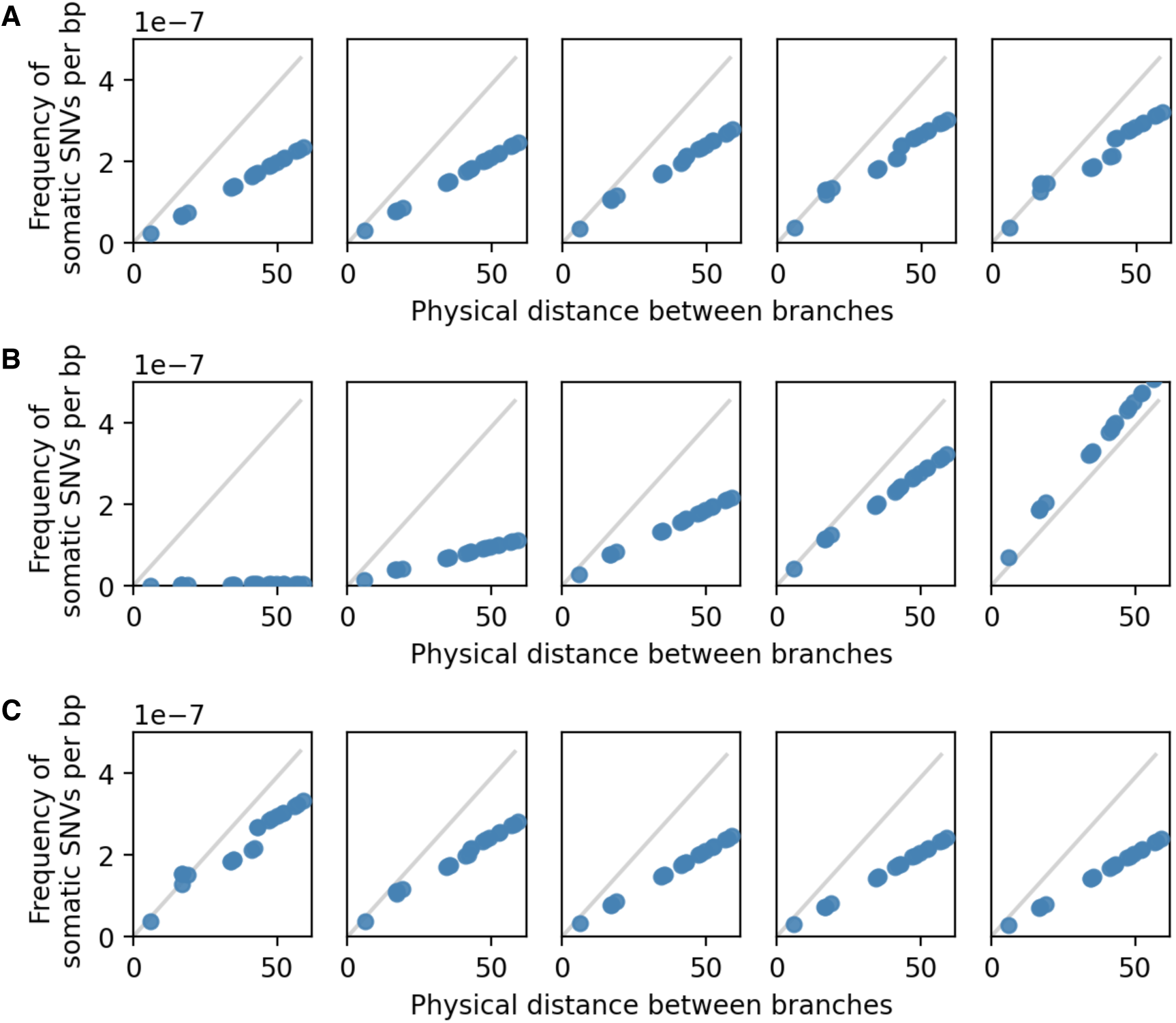
Parameter dependence of the model IV (replaced biased model). (A) Dependence on the number of stem cells. The values are 𝛼 = 1, 5, 15, 25, and 35, corresponding to the order of the subfigures from left to right. (B) Dependence on mutation rate per unit distance growth. The values are 𝑢 = 10^-10^, 1.7×10^-9^, 3.3×10^-9^, 4.9×10^-9^, and 8.0×10^-9^. (C) Dependence on the number of cell divisions per meter. The values are 𝑟 = 1, 3, 13, 23, and 33. While varying one parameter, other parameters were fixed with 𝛼 = 10, 𝑢 = 4.0×10^-9^, and 𝑟 = 5. The values highlighting parameter dependency were selected. The grey line is shown for comparison and comes from the regression of sequence data.

**Figure S6.**
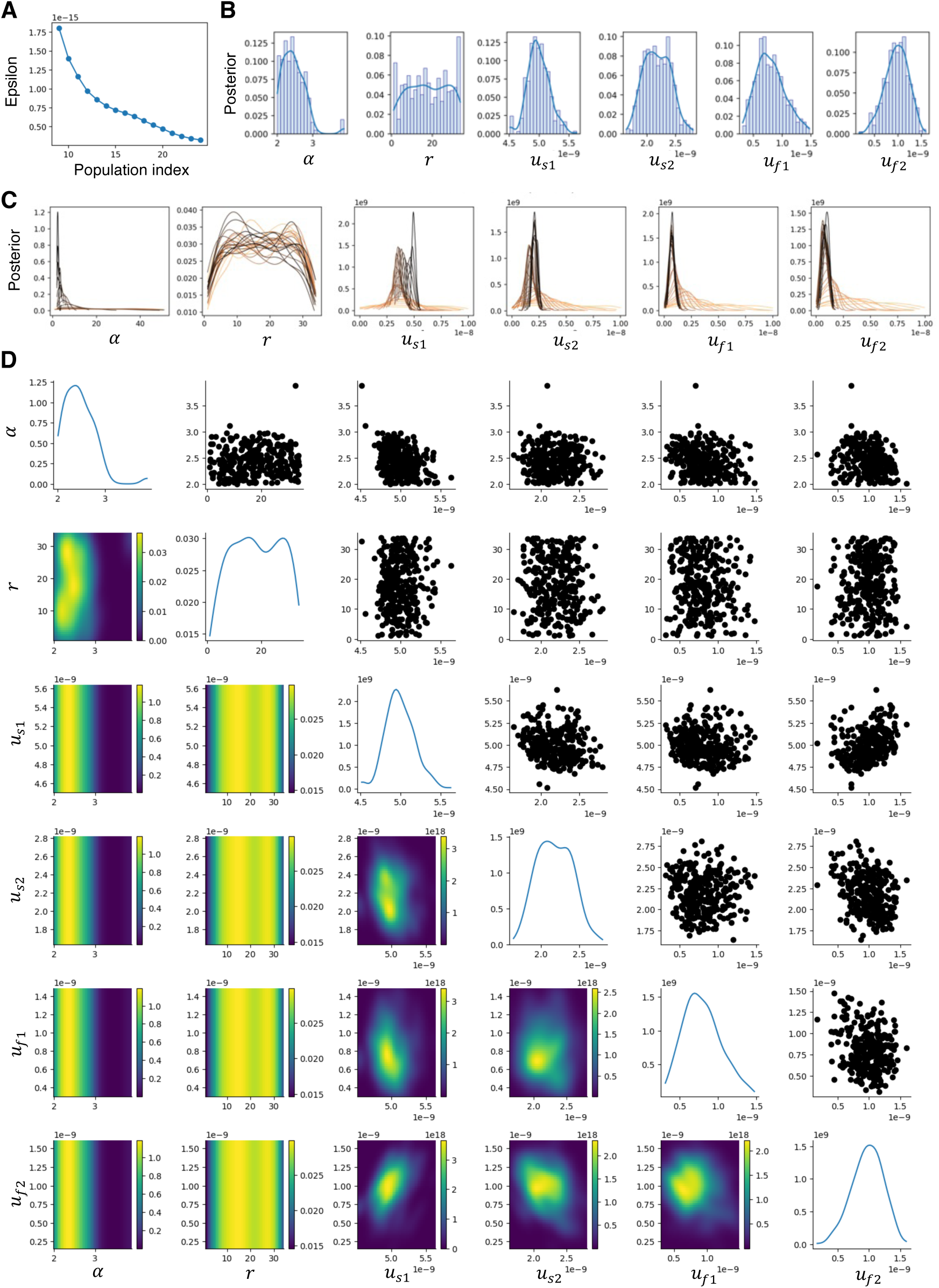
Results of parameter fitting for model I (conserved random model). (A) Epsilon, the minimum distance between model predictions and observations, over successive generations (population indices). From 10^th^ to the last generation are shown. (B) Posterior distributions of parameters at the final generation. Probability density distributions were estimated using kernel density estimation (KDE). (C) Posterior distributions over time. Colors indicate successive generations (population indices), with darker colors representing later generations. (D) Posterior distributions of parameters at the final generation and their pairwise relationships. Only KDE curves are shown.

**Figure S7.**
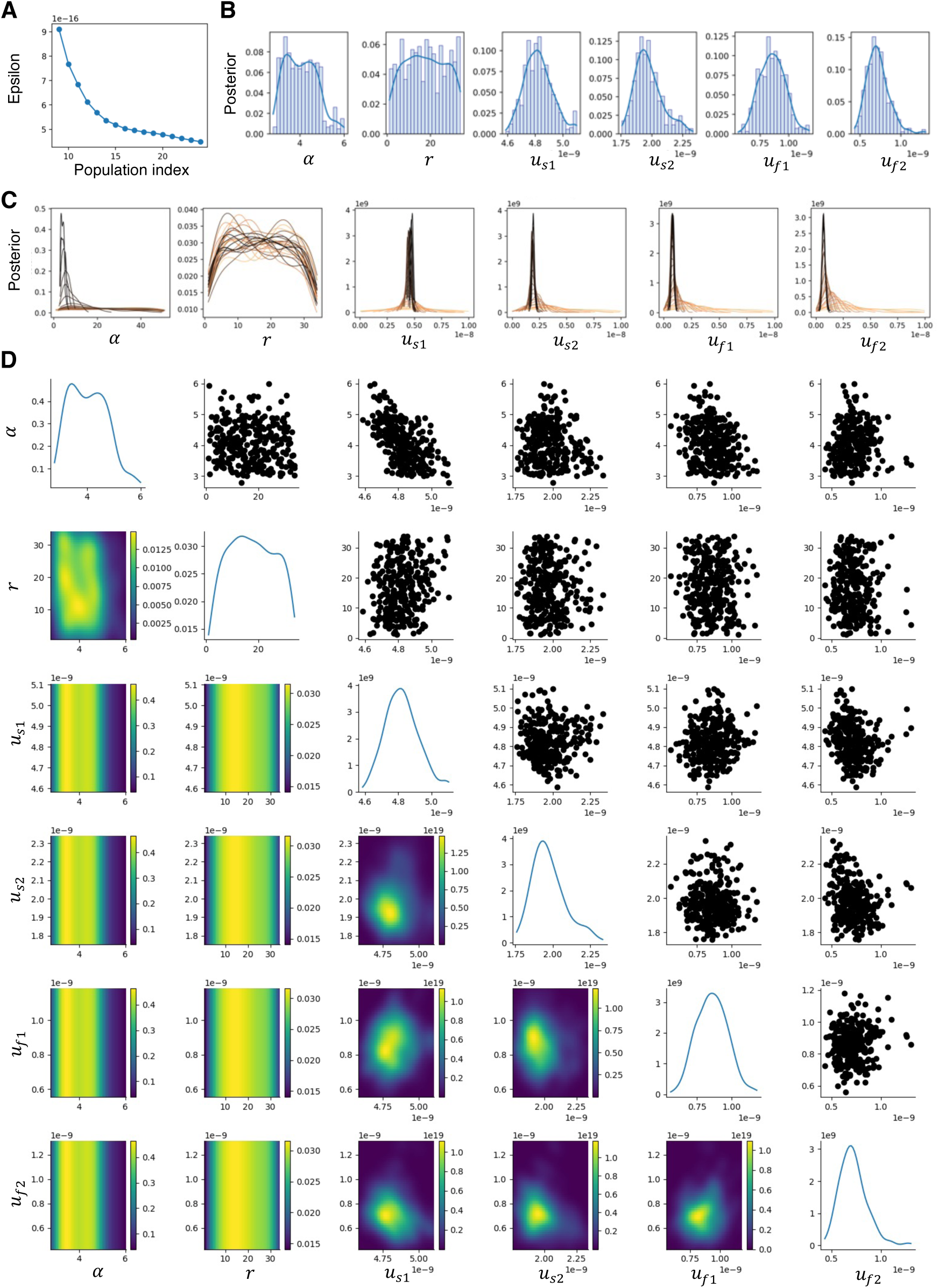
Results of parameter fitting for model II (conserved biased model). (A) Epsilon, the minimum distance between model predictions and observations, over successive generations (population indices). From 10^th^ to the last generation are shown. (B) Posterior distributions of parameters at the final generation. Probability density distributions were estimated using kernel density estimation (KDE). (C) Posterior distributions over time. Colors indicate successive generations (population indices), with darker colors representing later generations. (D) Posterior distributions of parameters at the final generation and their pairwise relationships. Only KDE curves are shown.

**Figure S8.**
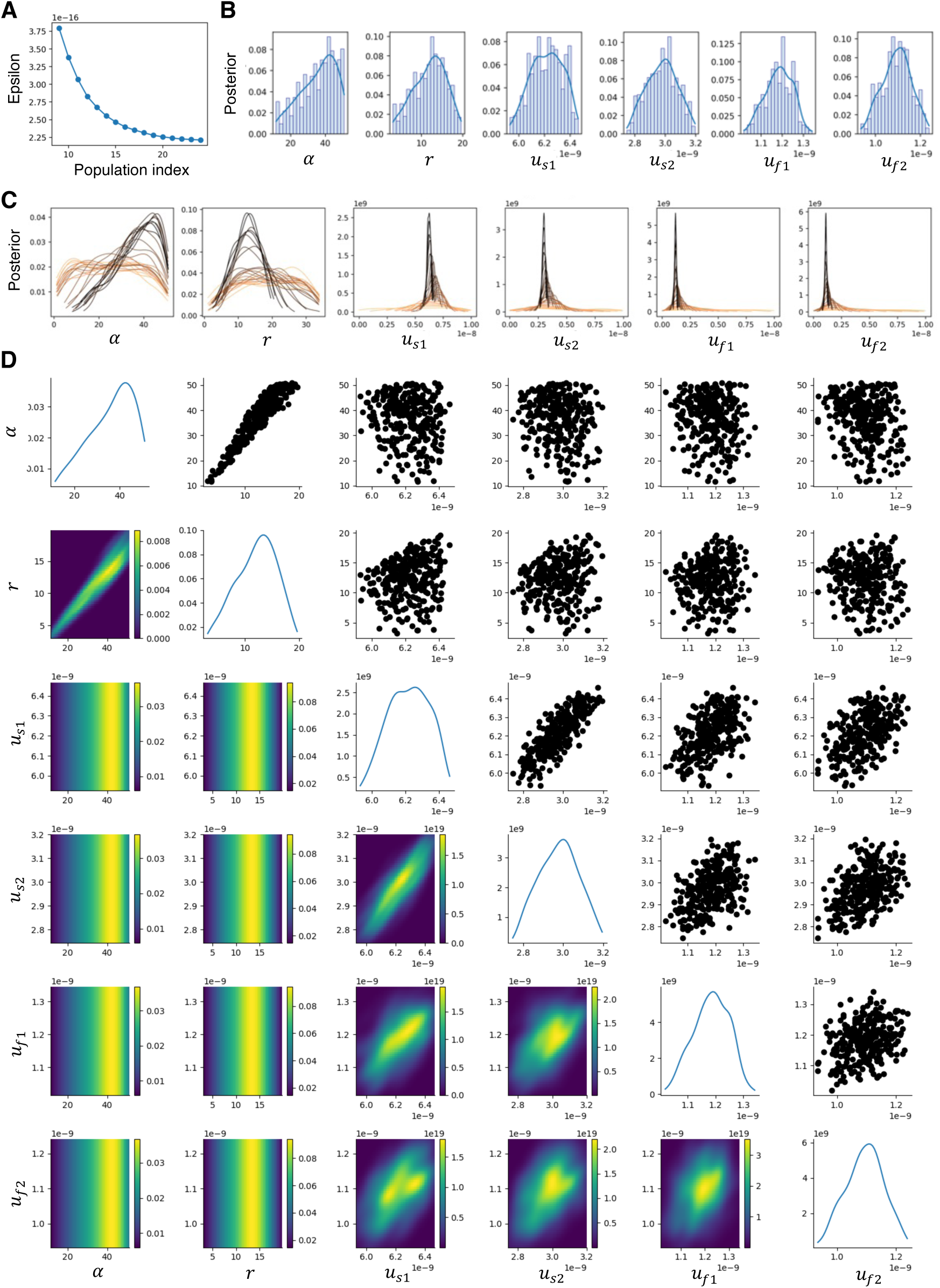
Results of parameter fitting for model III (replaced random model). (A) Epsilon, the minimum distance between model predictions and observations, over successive generations (population indices). From 10^th^ to the last generation are shown. (B) Posterior distributions of parameters at the final generation. Probability density distributions were estimated using kernel density estimation (KDE). (C) Posterior distributions over time. Colors indicate successive generations (population indices), with darker colors representing later generations. (D) Posterior distributions of parameters at the final generation and their pairwise relationships. Only KDE curves are shown.

**Figure S9.**
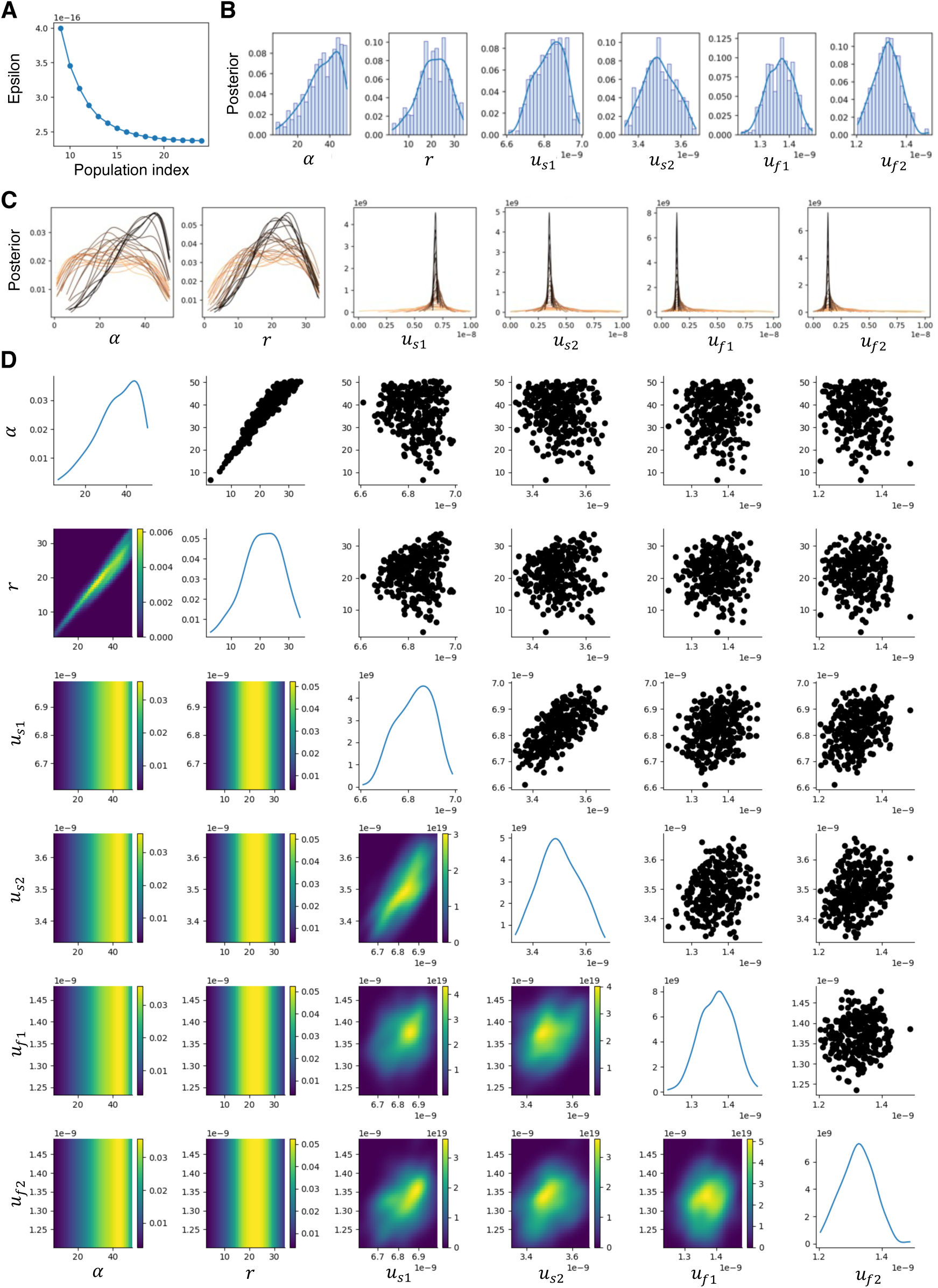
Results of parameter fitting for model IV (replaced biased model). (A) Epsilon, the minimum distance between model predictions and observations, over successive generations (population indices). From 10^th^ to the last generation are shown. (B) Posterior distributions of parameters at the final generation. Probability density distributions were estimated using kernel density estimation (KDE). (C) Posterior distributions over time. Colors indicate successive generations (population indices), with darker colors representing later generations. (D) Posterior distributions of parameters at the final generation and their pairwise relationships. Only KDE curves are shown.

**Figure S10.**
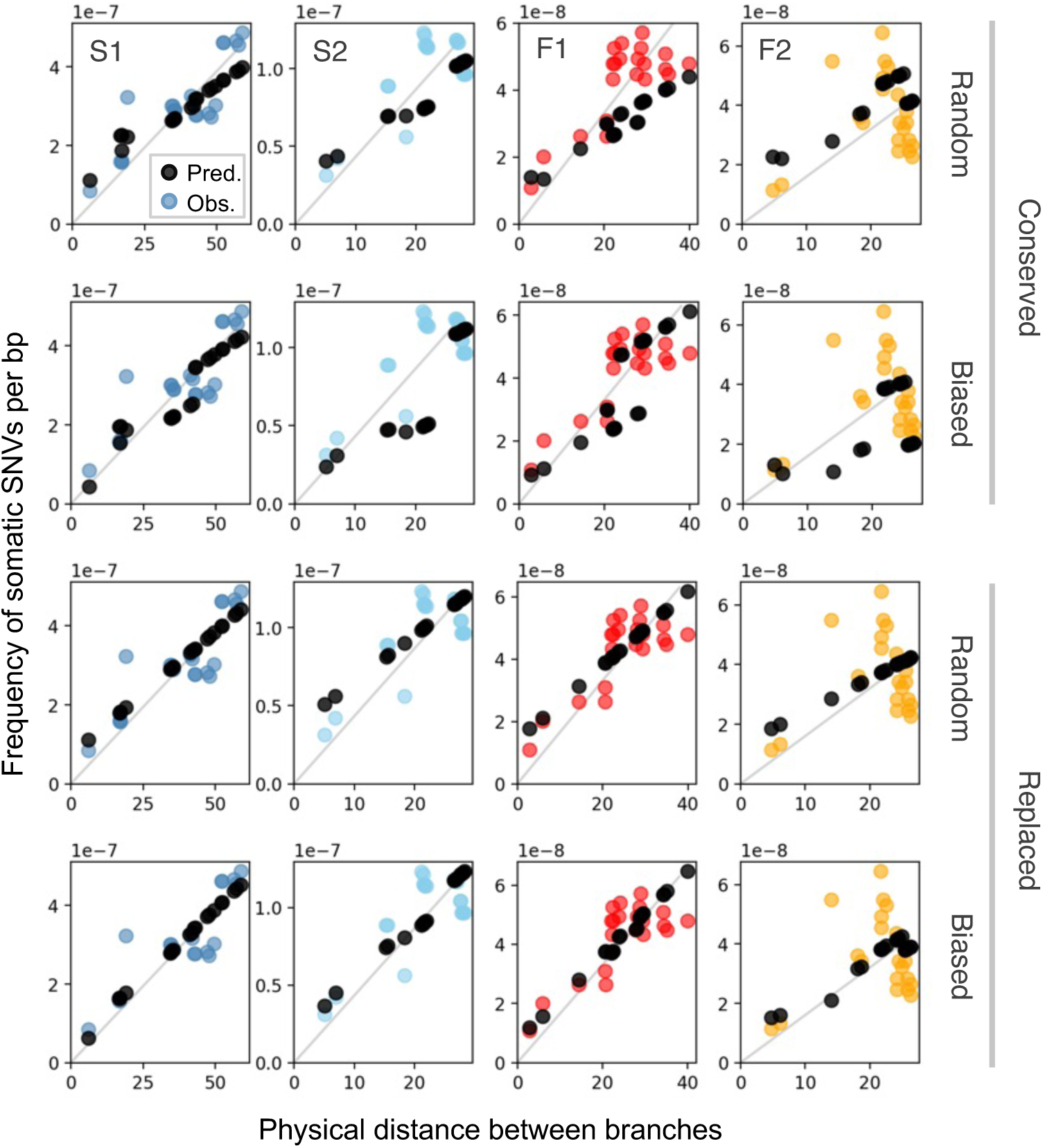
Inter-branch SNVs by the fitted model predictions, for all models (row) and individuals (column). The vertical axis represents the genetic difference between branch tips, i.e., the frequency of inter-branch SNVs per bp. The horizontal axis represents the physical distance between branch tips (m). Black circles indicate model predictions, while colored circles represent observations from different samples (S1: blue, S2: light blue, F1: red, F2: orange). The parameters used for predictions came from MAP estimates for each model and are listed in Table 1.

**Figure S11.**
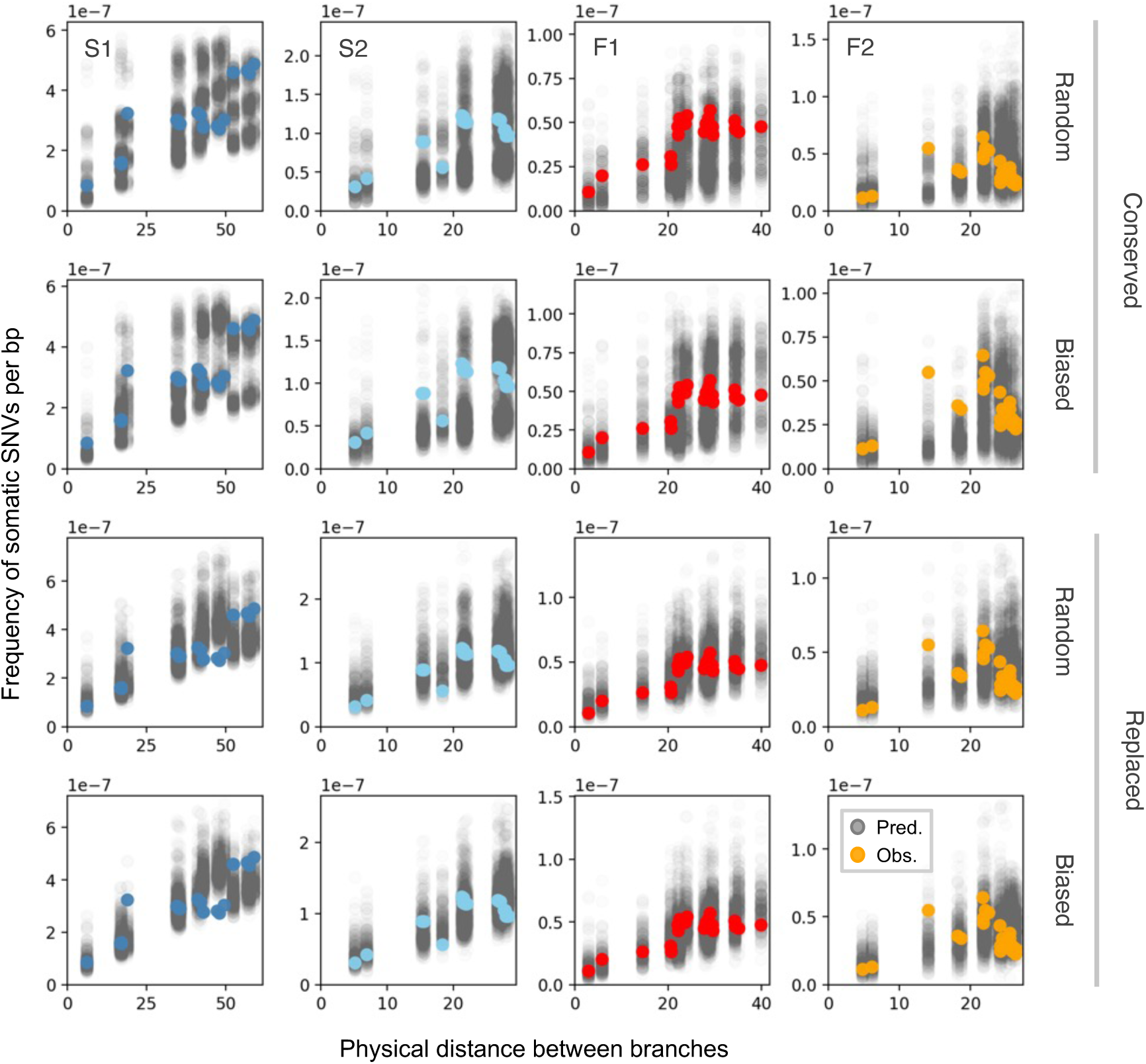
Simulated patterns of inter-branch SNVs. Inter-branch SNVs of predictions (grey points) vs observation (colored points). Predictions come from 250 runs of stochastic simulations using MAP estimates of parameters (Table 1).

**Figure S12.**
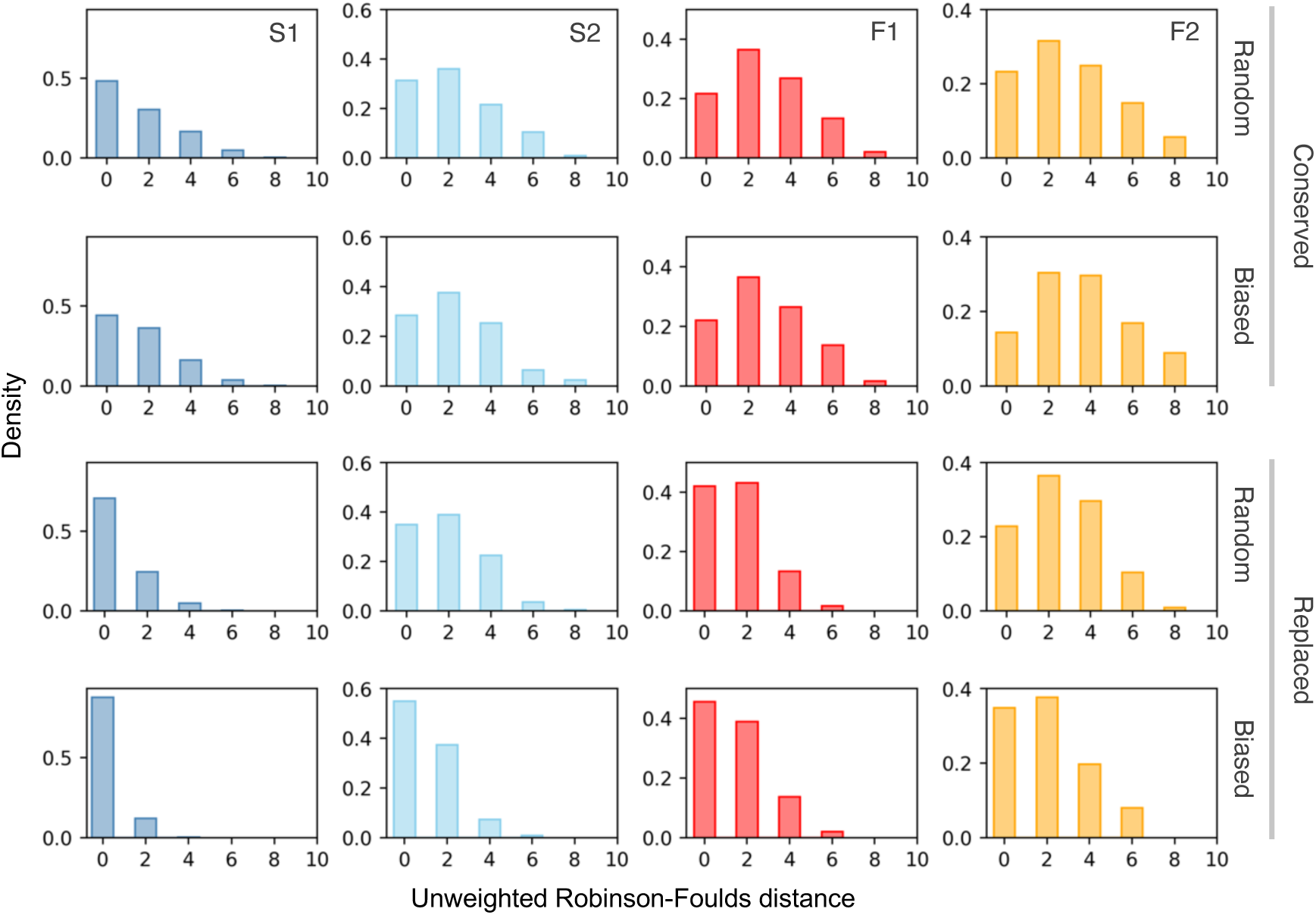
Distributions of unweighted Robinson-Foulds distances between physical architecture and somatic SNVs phylogeny predicted by the simulations. The distance quantifies the topological discordance, with 0 indicating perfect congruence. For all four individuals (S1, S2, F1, and F2; columns), the replaced models (bottom two rows) consistently produced a higher frequency of low distances compared to the conserved models (top two rows). This shift demonstrates that somatic drift is essential for generating genetic structures that align with the physical architecture. The distributions were derived from 250 runs of stochastic simulations for each model and individual.

**Figure S13.**
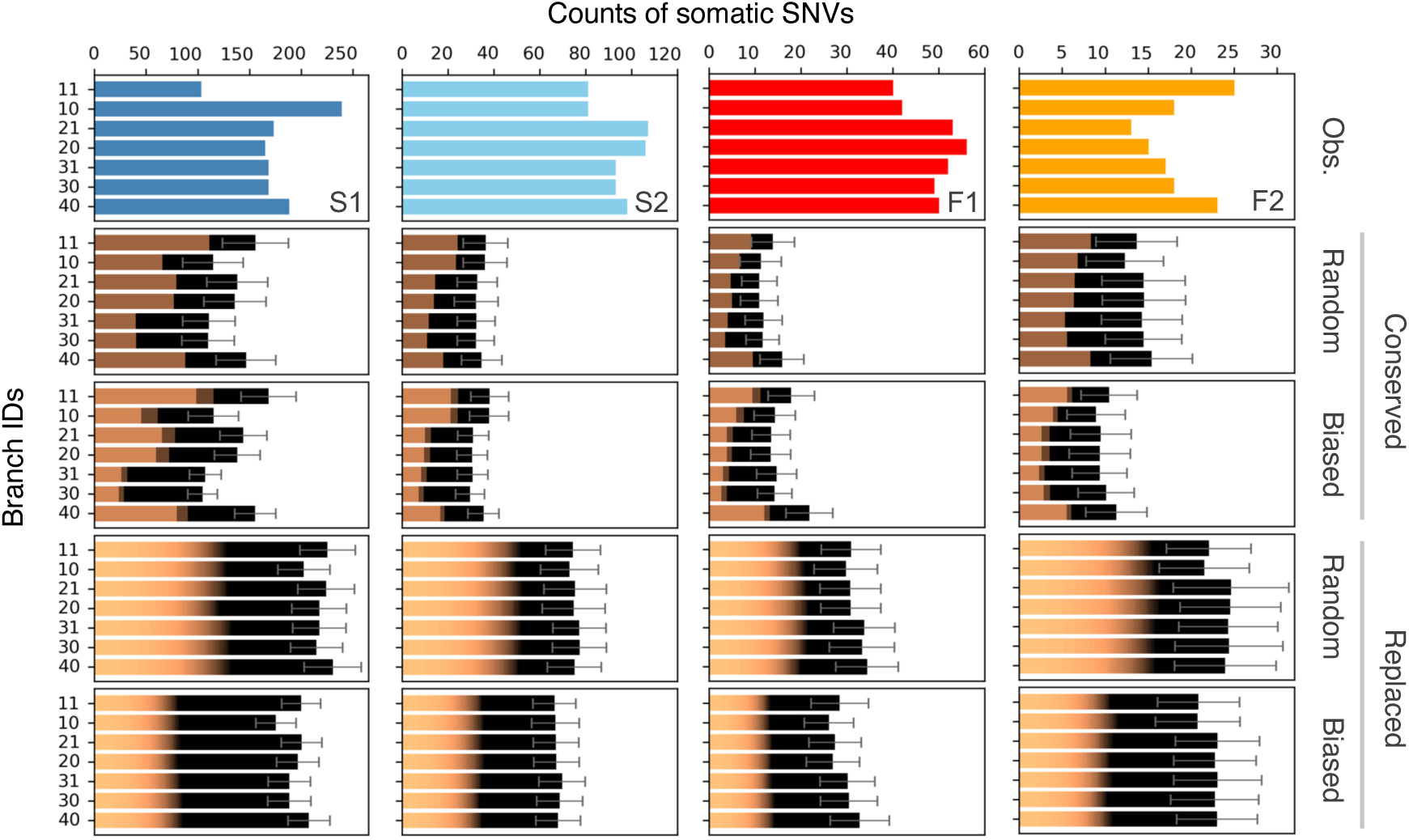
Number of accumulated SNVs at the tips of branches. Distribution of the distance were derived from 250 runs of simulations, for all models (row) and individuals (column). (B) Number of accumulated mutations at the tips of branches. Columns correspond to the tree individuals. Colors in predictions indicate the frequency of SNVs in a SAM (darker the higher). Black represents fixed SNVs.

**Figure S14.**
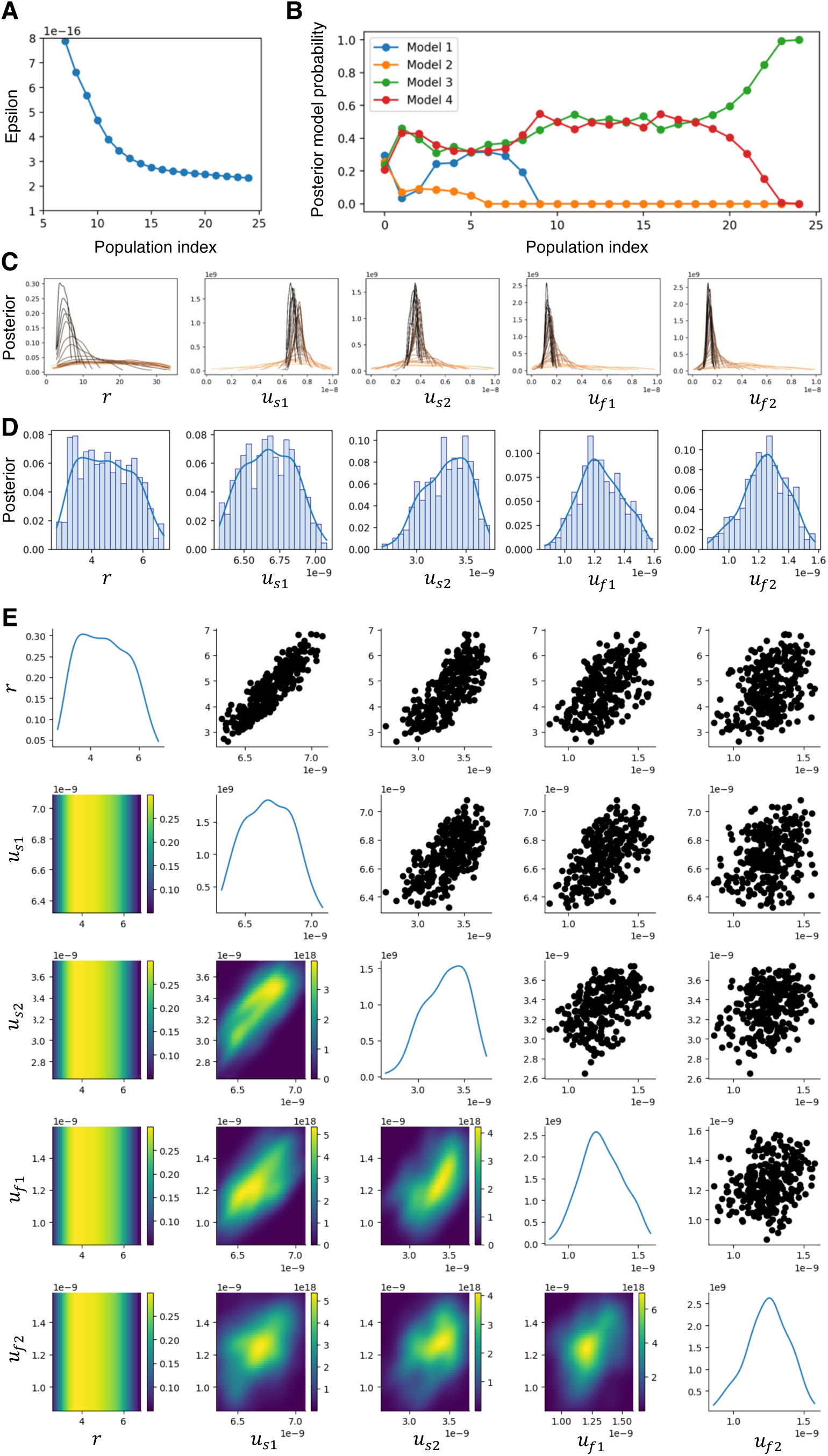
Results of model fitting under. 𝜶 = 𝟏𝟎. (A) Epsilon, the minimum distance between model predictions and observations, over successive generations (population indices). (B) Posterior model probabilities over generations. Model 3 was selected as the best-supported model. Model 1: conserved random, Model 2: conserved biased, Model 3: replaced random, Model 4: replaced biased. (C) Posterior distributions over time for the selected model. Colors indicate successive generations (population indices): darker colors correspond to later generations. Posterior distributions of parameters at the final generation (D) and their pairwise relationships (E). Distributions were estimated using kernel density estimation.

